# Lipid droplet protein Perilipin 2 is critical for the regulation of insulin secretion through beta cell lipophagy and glucagon expression in pancreatic islets

**DOI:** 10.1101/2024.11.17.624030

**Authors:** Siming Liu, Israel Wipf, Aditya Joglekar, Aidan Freshly, Corinne E Bovee, Lucy Kim, Syreine L Richtsmeier, Spencer Peachee, Shayla Kopriva, Anamika Vikram, Dalal El Ladiki, Fatma Ilerisoy, Beyza Ilerisoy, Gianna Sagona, Claudia Jun, Michelle Giedt, Tina L Tootle, James Ankrum, Yumi Imai

## Abstract

Knockdown (KD) of lipid droplet (LD) protein perilipin 2 (PLIN2) in beta cells impairs glucose-stimulated insulin secretion (GSIS) and mitochondrial function. Here, we addressed a pathway responsible for compromised mitochondrial integrity in PLIN2 KD beta cells. In PLIN2 KD human islets, mitochondria were fragmented in beta cells but not in alpha cells. Glucagon but not insulin level was elevated. While the formation of early LDs followed by fluorescent fatty acids (FA) analog Bodipy C12 (C12) was preserved, C12 accumulated in mitochondria over time in PLIN2 KD INS-1 cells. A lysosomal acid lipase inhibitor Lali2 prevented C12 transfer to mitochondria, mitochondrial fragmentation, and the impairment of GSIS. Direct interactions between LD-lysosome and lysosome-mitochondria were increased in PLIN2 KD INS-1 cells. Thus, FA released from LDs by microlipophagy cause mitochondrial changes and impair GSIS in PLIN2 KD beta cells. Interestingly, glucolipotoxic condition (GLT) caused C12 accumulation and mitochondrial fragmentation similar to PLIN2 KD in beta cells. Moreover, Lali2 reversed mitochondrial fragmentation and improved GSIS in human islets under GLT. In summary, PLIN2 regulates microlipophagy to prevent excess FA flux to mitochondria in beta cells. This pathway also contributes to GSIS impairment when LD pool expands under nutrient load in beta cells.

---

Beta cell dysfunction is the key pathology of type 2 diabetes (T2D).^1^ Functional decline of beta cells in T2D occurs early from prediabetes stage and is progressive in nature contributing to the need for insulin therapy in subjects with long-term T2D.^2^ Nutrient overload is a major risk factor for T2D and can provoke stress responses known to drive beta cell demise such as inflammation, endoplasmic reticulum (ER) stress, and mitochondrial dysfunction.^3^ Multiple studies have indicated that a strong association between hyperlipidemia (fatty acids (FA), triglycerides (TG), ceramides, acyl-carnitines) and development of T2D. ^4–7^ In addition to circulating lipids, pancreatic islets can be exposed to lipids from surrounding adipocytes in pancreas. Indeed, the correlation between pancreatic fat and beta cell dysfunction has been reported.^8^ Unlike glucose uptake that is regulated by glucose transporters, plasma membrane is more permissive to FA and the elevation of environmental FA is sufficient to promote FA influx to cells.^9^ Thus, an intracellular mechanism to regulate trafficking of free FA to minimize cellular level of free FA is critical considering toxic nature of free FA.^10,11^

A lipid droplet (LD) is an organelle that serves as a hub of intracellular lipid metabolism with its ability to store neutral lipids and release lipids in a temporally and spatially regulated manner.^12^ In addition, LD scavenges cytotoxic FA metabolites and serves as the first defender to maintain lipid homeostasis within a cell.^13^ Five members of perilipin family of protein (PLIN1-5), each with a unique expression pattern, reside on the surface of LDs and plays a critical role in LD formation and degradation in a cell type specific manner.^14,15^ Pancreatic beta cells express PLIN2 predominantly in a manner that positively correlates with TG levels within pancreatic islets and beta cells.^9,16^ Overexpression of PLIN2 is sufficient to expand TG pool without impairing insulin secretion in MIN6 cells.^16^ Although the improvement in beta cell survival was reported by the whole body KO of PLIN2 in Akita mice,^17^ downregulation of PLIN2 impairs insulin secretion in MIN6 cells, INS-1 cells, beta cell specific knockout (KO) mice, EndoC cells, and human pseudoislets indicating that PLIN2 is critical for maintaining normal insulin secretion^16,18,19^ However, it has not been clear why PLIN2 deficiency in beta cells impairs insulin secretion. Interestingly, PLIN2 KO or knockdown (KD) reduced OXPHOS proteins in INS-1 cells, mouse islets, and human pseudoislets and lowered oxygen consumption rate (OCR) in INS-1 cells and mouse islets. Thus, PLIN2 deficiency appears to compromise the integrity of mitochondrial in beta cells.^18^ We also noted that a fluorescent FA analog Bodipy 558/568 C12 (Bodipy C12)^20^ distributes to mitochondria instead of LDs when added to PLIN2 KD INS-1 cells.^18^

In this study, we aim to determine a pathway by which PLIN2 deficiency leads to Bodipy C12 accumulation in mitochondria, impaired mitochondrial integrity, and lower glucose-stimulated insulin secretion (GSIS). We also addressed whether PLIN2 deficiency impacts human beta cells and alpha cells in a similar manner. We found that PLIN2 deficiency increases LD-lysosome and lysosome-mitochondria interactions and that the inhibition of TG degradation in lysosome prevents Bodipy C12 accumulation in mitochondria, mitochondrial fragmentation, and beta cell dysfunction. Although PLIN2 KD reduced LD size in human alpha cells, a morphological change of mitochondria was not seen in alpha cells and glucagon secretion was not impaired. The pathway of FA trafficking from LDs to mitochondria identified in PLIN2 KD beta cells appears to be operative under nutrient stress as well. The inhibition of lysosomal acid lipase (LIPA) prevented mitochondrial fragmentation and improved GSIS in human beta cells exposed to nutrient load.

## RESULTS

### PLIN2 downregulation has distinct effects on hormone production and secretion in human beta and alpha cells

We previously reported that PLIN2 KD in human pseudoislets impairs insulin secretion.^18^ Considering that *PLIN2* is also expressed in alpha cells, we addressed whether PLIN2 downregulation affects glucagon secretion.^21^ In agreement with previous data,^18^ there was no difference in *INS* expression between scramble control (Cont) and PLIN2 KD human pseudoislets when tested in 12 donor islets. With a higher number of donors, *GCG* expression reached a statistically significant 2.1-fold increase in PLIN2 KD human pseudoislets (Fig. 1A). The change in gene expression was not associated with an increase in the proportion of alpha or beta cells defined by the reactivity to anti-GCG and INS antibodies in human islet cells after PLIN2 KD (Fig. 1B). The proportion of cells negative for both INS and GCG did not change either (data not shown). Reproducing our previous data,^18^ insulin secretion in response to both glucose and KCl was reduced in PLIN2 KD pseudoislets without a change in INS contents (Fig. 1C, D). GCG contents were significantly elevated to 2.5-fold of Cont in PLIN2 KD pseudoislets indicating that GCG is increased at a peptide level as well (Fig. 1E). Glucagon secretion showed a trend of increase in PLIN2KD pseudoislets but did not reach statistical significance compared with Cont treated with the same concentration of secretagogues (Fig. 1F). Also, the suppression of glucagon secretion by 7 mM glucose and the stimulation by KCl were seen in both Cont and PLIN2 KD pseudoislets indicating that the regulation of glucagon secretion is grossly present in PLIN2 KD pseudoislets.

**Figure 1.**
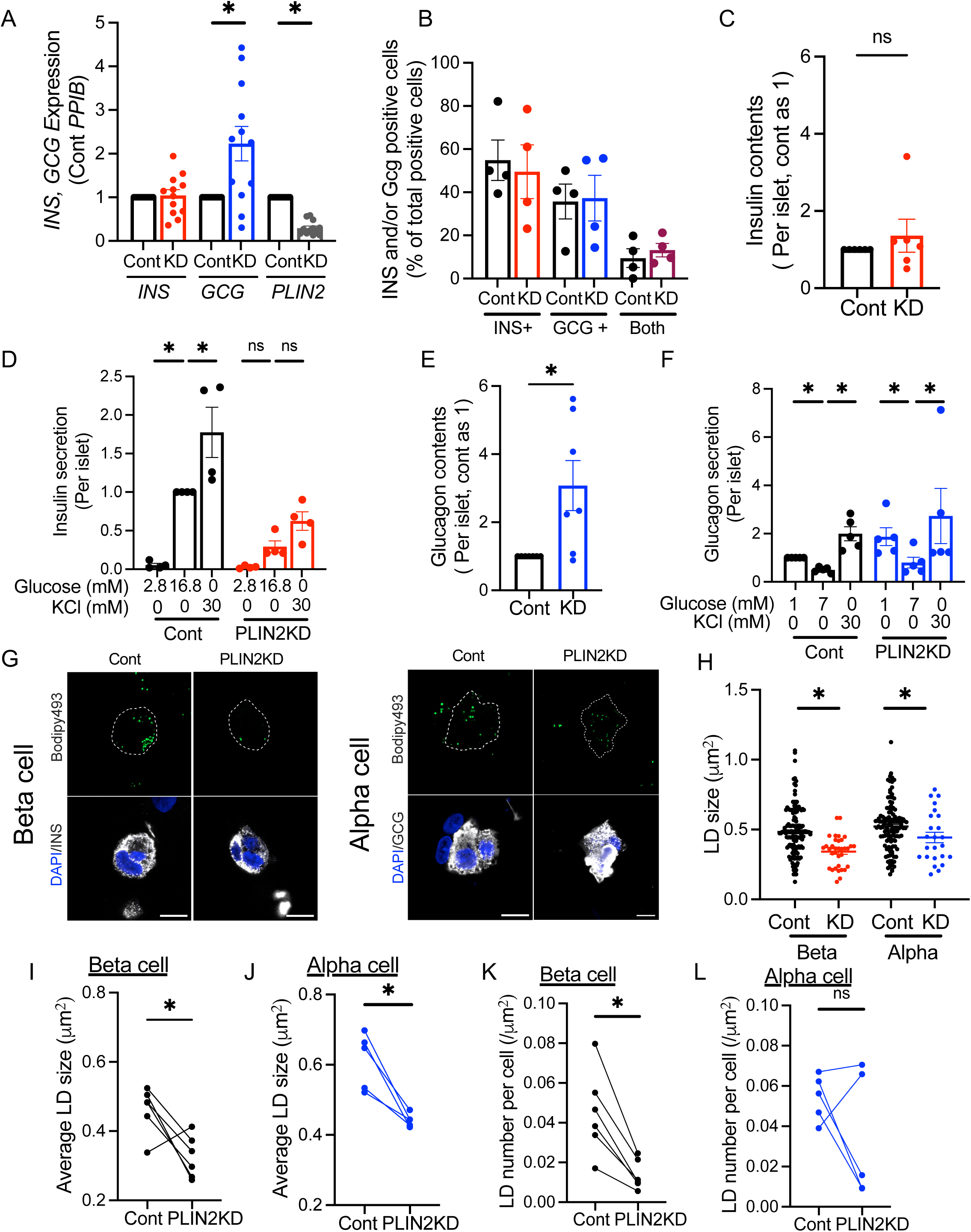
human PLIN2 downregulation impairs insulin secretion while increasing glucagon in human islets. PLIN2 was downregulated in human islet cells by lentivirus expressing shPLIN2 (PLIN2KD) using scramble shRNA as control (Cont). Either human pseudoislets were created or cells were plated on a confocal dish as in methods. (A) RT-qPCR determined expression of indicated genes in human pseudoislets with (KD) or without (Cont) downregulation of PLIN2. Value was expressed taking the expression of Cont as 1 for each donor. n= 12 donor islets. (B) Proportion of insulin positive (INS), glucagon positive (GCG) or INS/GCG double positive cells (both) were determined by immunofluorescence imaging of human islet cells. n=4 donor islets. (C) INS contents per islet was compared between Cont and PLIN2KD pseudoislets and expressed taking Cont as 1. n=6 donors. (D) INS secretion from Cont and PLIN2KD pseudoislets in response to indicated concentration of glucose or KCl was corrected per islet and expressed taking INS secretion at 16.8 mM glucose in Cont islets as 1. n=4 donors. (E) GCG contents per islet was compared between Cont and PLIN2KD pseudoislets and expressed taking Cont as 1. n=7 donors. (F) GCG secretion from Cont and PLIN2KD pseudoislets in response to indicated concentration of glucose or KCl was corrected per islet and expressed taking GCG secretion at 1 mM glucose in Cont islets as 1. n=5 donors. (G-L) Representative images and morphometry of lipid droplets (LDs) in Cont and PLIN2 KD human alpha and beta cells by confocal microscopy. (G) LDs were visualized by Bodipy 493/503 (Bodipy 493, green), beta cells by anti-INS antibody, and alpha cells by anti-GCG antibody. Cell border in a Bodipy 493 image is indicated by dotted line. Scale bar, 10 μm. (H) Size distribution of LDs in beta and alpha cells in a representative donor. (I, J) Average size and (K, L) number of LD per cell area in beta cells (I, K, n=6 donors) and alpha cells (J, L, n=5 donors). A dot represents each donor and the number from the same donor is connected by line. Data are mean ± SEM. (D, F), *; p<0.05 by One-way ANOVA test. The rest, *; p<0.05 by Student’s t-test.

With differential effects of PLIN2 deficiency on hormone contents and secretion between INS and GCG, we compared LD morphometry between human beta and alpha cells plated on a type IV collagen coated confocal dish, a condition that maintains the differentiated status of human beta cells, GSIS, and dynamic changes of LDs and mitochondria.^22,23^ PLIN2 KD reduced the size of LDs in both beta and alpha cells indicating that PLIN2 determines LD size in both cells (Fig. 1G-J). However, the reduction in the number of LDs was more prominent in beta cells than alpha cells, which could be due to high expression of another lipid droplet protein PLIN3 in alpha cells (Fig. 1K,L).^21^ PLIN3 is known as a lipid droplet protein that is primarily recruited to small size LDs and known to compensate for the loss of PLIN2.^24,25^

### PLIN2 KD fragments mitochondria of human beta but not that of alpha cells

When PLIN2 deficiency dysregulates insulin secretion in mouse islets, INS-1 cells, and human pseudoislets, we previously observed the reduction of OXHPOS proteins in all three models, low oxygen consumption rate (OCR) in INS-1 cells and mouse islets, and fragmented mitochondria in INS-1 cells (Fig. 2A).^18^ Thus, the reduced integrity of mitochondria is a possible culprit behind impaired insulin secretion in PLIN2 KD beta cells. Therefore, we tested whether PLIN2 KD also fragments mitochondria in human islet cells. Human beta cells treated with shPLIN2 showed reduction in form factor, mitochondrial length, and aspect ratio compatible with more fragmented morphology of mitochondria (Fig. 2B-E, supplementary Fig. 1A-D). In comparison, PLIN2 KD did not fragment mitochondria in alpha cells indicating that beta cell mitochondria are more affected by PLIN2 KD than alpha cells (Fig. 2F-H). Therefore, we focused our further study to determine a mechanism by which PLIN2 KD causes mitochondrial fragmentation and impairs insulin secretion in beta cells.

**Figure 2.**
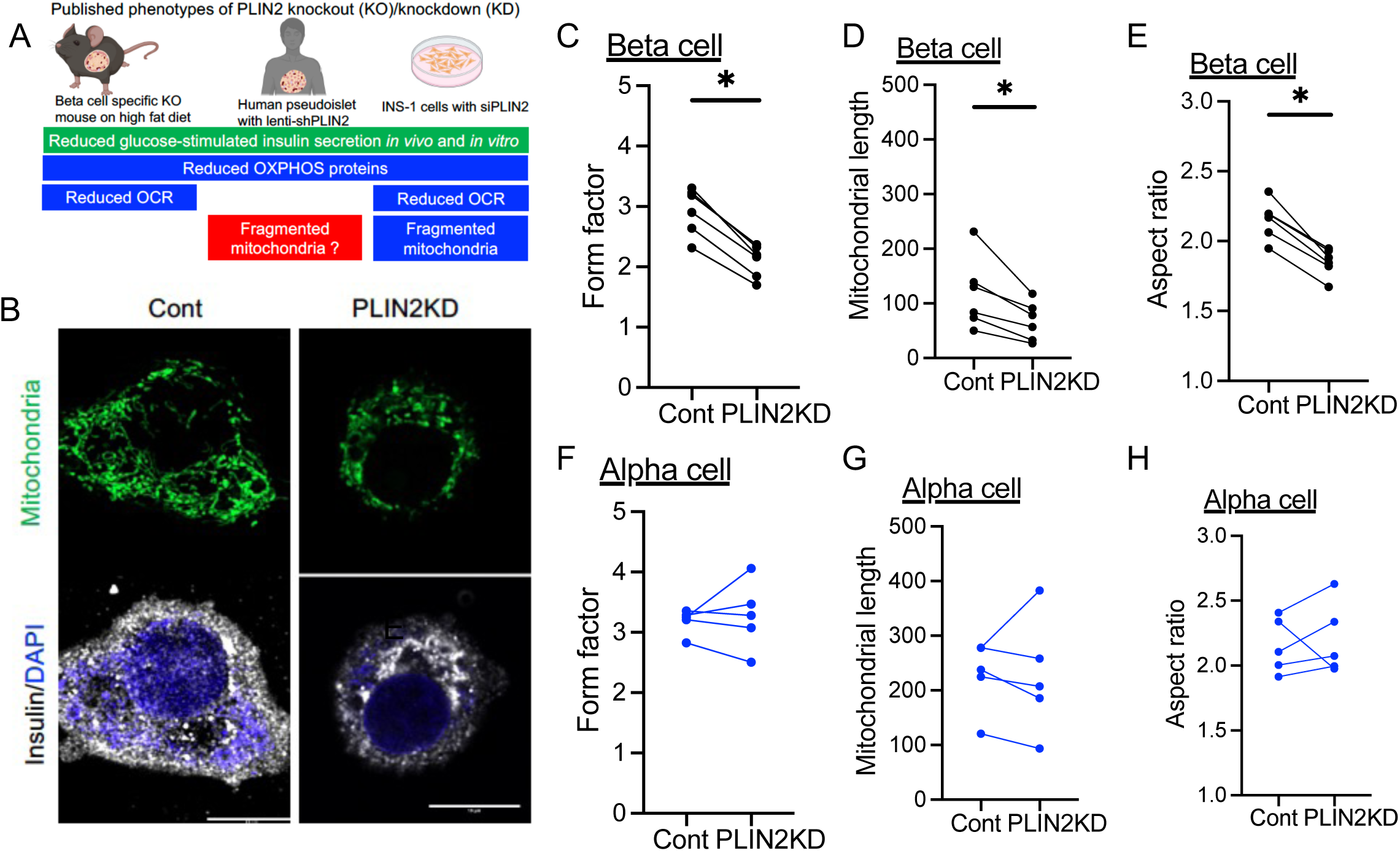
human PLIN2 downregulation fragments mitochondria in human beta cells, but not in alpha cells. (A) Phenotypes of PLIN2 downregulation in three beta cell models published in ^18^. We now address whether mitochondria fragmentation is seen in human beta and alpha cells in (B-H). PLIN2 was downregulated in human islet cells plate on a confocal dish by lentivirus expressing shPLIN2 (PLIN2KD) using scramble shRNA as control (Cont). (B-H) The morphology of mitochondria was assessed in (B-E) beta cells and (F-H) alpha cells. (B) Representative images of beta cells marked by anti-insulin antibody (white). Mitochondria was visualized by mitochondria-GFP (green) and nuclei by DAPI (blue). Scale bar, 10 μm. (C, F) Form factors, (D, G) mitochondrial length, and (E, H) aspect ratio as parameters of mitochondria shape. n= 5-6 donors. A dot represents each donor and the number from the same donor is connected by line. Data are mean ± SEM. *; p<0.05 by Student’s t-test.

### The formation of nascent LDs is maintained in PLIN2 KD INS-1 cells

Nutrient load is known to increase the fragmentation of mitochondria in beta cells.^23,26^ When cells are metabolically labeled with a fluorescent FA analog Bodipy C12 overnight, Bodipy C12 localized in mitochondria instead of LDs in PLIN2 KD INS-1 cells implicating that PLIN2 KD may overload mitochondria with FA and fragment mitochondria.^18^ To address a mechanism responsible for the increased Bodipy C12 localization in mitochondria, we monitored time course of Bodipy C12 distribution in Cont and PLIN2 KD INS-1 cells (Fig. 3A). LDs are formed at ER by budding off neutral lipids accumulated in an outer layer of ER membrane creating a nascent LD particle consisting of a lipid core covered by a monolayer of phospholipids.^27^ When the formation of nascent LDs is quantified by a short time labeling of cells with Bodipy C12, the size of nascent LDs did not differ at 30 min but started to decline in PLIN2 KD INS-1 cells at 60 min of metabolic labeling with Bodipy C12 (Fig. 3B-C). At these early time points, the number of nascent LDs corrected per cell area did not differ between control and PLIN2 KD INS-1 cells indicating that the formation of nascent LDs is not impaired in PLIN2 KD INS-1 cells (Fig. 3D). Bodipy C12 did not colocalize with mitochondria significantly at 1 h after the addition of Bodipy C12 in either Cont or PLIN2 KD cells (Fig. 3E, F). By 6 h, Bodipy C12 was condensed in LDs represented by bright dots in Cont INS-1 cells (Fig. 3E). In contrast, PLIN2KD cells had little LDs and Bodipy C12 showed a reticular pattern of distribution with high colocalization with mitochondria (Fig. 3F). Therefore, the accumulation of Bodipy C12 in mitochondria is a later event compared with the formation of nascent LDs that is not impaired in PLIN2 KD INS-1 cells. Data collectively point that Bodipy C12 trafficking to mitochondria occurs after degradation of LDs that are initially formed.

**Figure 3.**
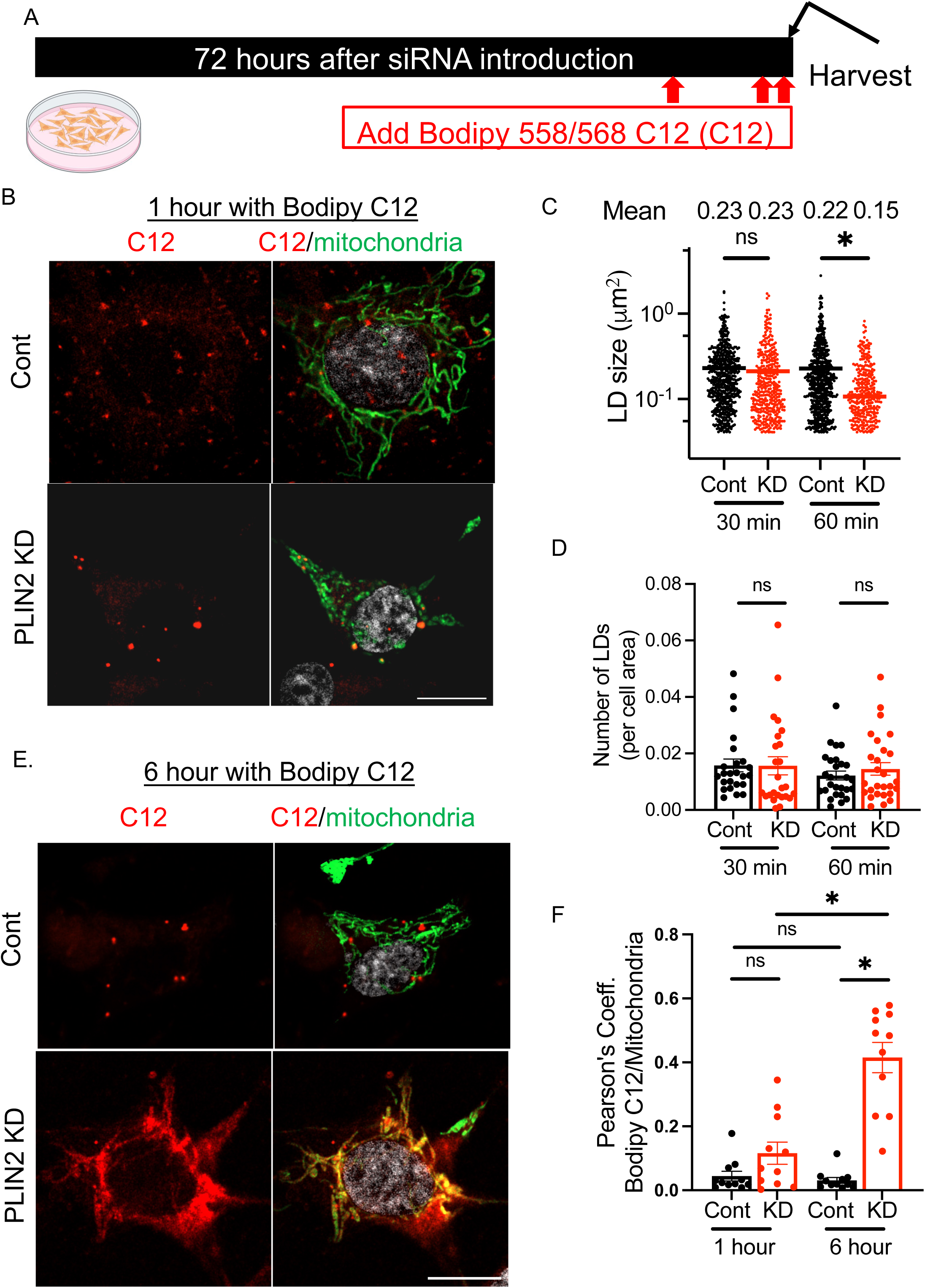
The production of nascent lipid droplets is not reduced in PLIN2 downregulated INS-1 cells. (A) Experimental design showing that control (Cont) and PLIN2 deficient (PLIN2KD) INS-1 cells were metabolically labeled with fluorescent FA analog Bodipy 558/568 C12 (C12) for 0.5 to 6 h and harvested for confocal microscopy. (B) Representative images showing C12 signal and mitochondria visualized by GFP in green after 1 h incubation with C12. (C) Size and (D) number of nascent lipid droplets (LDs) identified as C12 marked punctate at the indicated time after incubation with C12. (C) Each dot represents an individual LD. n= 215-472 combining three independent experiments. Y axis is log 10 due to wide distribution of data. (D) Each dot represents the number of LDs in an individual image corrected for cell area. n= 24-27 images from three independent experiments. (E) Representative images showing C12 signal and mitochondria visualized by GFP in green after 6 h incubation with C12. (F) Pearson’s coefficient of C12 and mitochondria GFP. n=11 images, representative of three experiments. All scale bars are 10 μm. ns; not significant, *; p<0.05 by Sidak’s multiple comparison test.

### The inhibition of lysosomal acid lipase prevents Bodipy C12 distribution to mitochondria in PLIN2 KD INS-1 cells

Degradation of neutral lipids in LDs can be catalyzed by two TG lipases, one being PNPLA2 (also known as ATGL) that acts on the surface of LDs and the other being lysosomal acid lipase (LIPA) that degrades TG after delivery to lysosome.^28^ Both lipases are expressed and reported to hydrolyze TG in beta cells.^29,30^ Thus, we tested whether the inhibition of PNPLA2 and LIPA restore lipid depots and TG levels in PLIN2 KD INS-1 cells, which appears to have accelerated degradation of LDs. In Cont cells, the inhibition of PNPLA2 by atglistatin (ATGLi) did not increase the number of Bodipy C12 positive lipid depots significantly (Fig. 4A, B) but increased the size of each lipid depot (Fig. 4C), a pattern expected for PNPLA2 inhibition that occurs on the surface of LDs. In comparison, LIPA inhibitor lalistat 2 (Lali2) increased the number of lipid depots dramatically (Fig. 4B). Although large size lipid depots increased mildly, the increase in small size lipid depots was more prominent in Lali2 treated control INS-1 cells making overall size distribution not significantly different from DMSO treated control (Fig. 4C). The predominant increase of small lipid depots after Lali2 treatment is compatible with the accumulation of TG in lysosomes. Thus, each inhibitor showed changes in lipid depot reflective of its site of action in Cont INS-1 cells. PLIN2 KD INS-1 cells in the absences of lipase inhibitors showed lower number and size of lipid depots compared with Cont cells in agreement with human beta cells (Fig. 1G-J, Fig. 4A-C). ATGLi increased size of lipid depots in PLIN2 KD INS-1 cells indicating that PNPLA2 contributes to degradation of LDs in PLIN2 KD INS-1 cells to some extent (Fig. 4C). However, Lali2 caused a much drastic increase in both size and number of lipid depots in PLIN2 KD INS-1 cells (Fig. 4B, C). Lali2 treatment significantly raised TG contents in control INS-1 cells indicating that LIPA actively degrades LD under nutritionally sufficient condition in INS-1 cells (Fig. 4D). TG was reduced in PLIN2 KD INS-1 cells as expected but increased significantly in the presence of Lali2 providing a biochemical proof that lipophagy of TG rich LD is active in PLIN2 KD INS-1 cells. More importantly, the prevention of lysosomal degradation of TG by Lali2 was sufficient to reduce Bodipy C12 in mitochondria in PLIN2 KD INS-1 cells implicating that Bodipy C12 travels to mitochondria after lysosomal degradation of LDs (Fig. 4E, F). Due to high autofluorescence and the presence of lipofuscin that have wide emission and excitation, Pearson’s coefficient could not be determined in human beta cells.^31,32^

**Figure 4.**
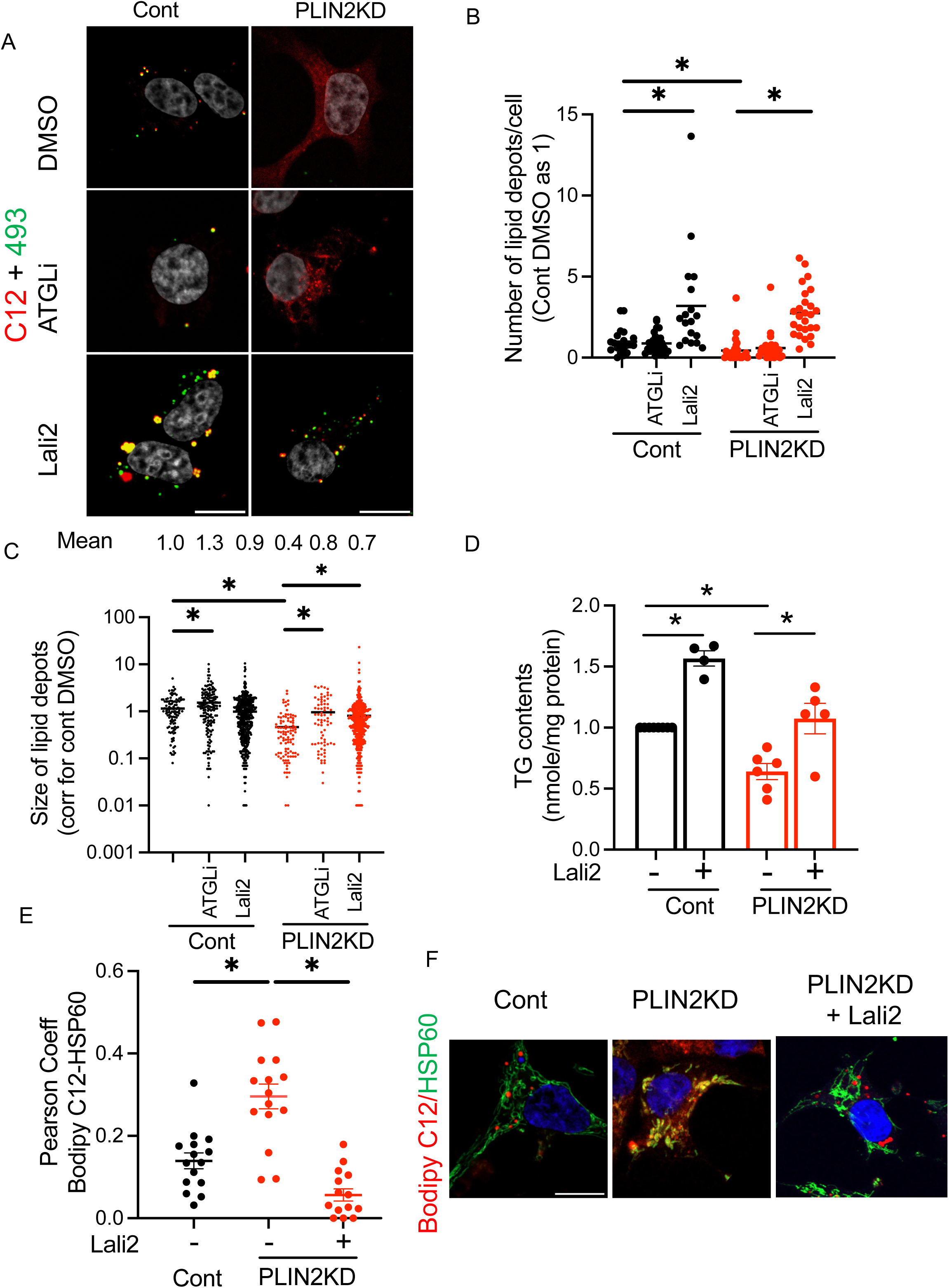
A LIPA inhibitor increases neutral lipid depots and prevents Bodipy C12 localization to mitochondria in PLIN2 downregulated INS-1 cells. (A) Representative confocal images of control (Cont) and PLIN2 deficient (PLIN2KD) INS-1 cells metabolically labeled with Bodipy C12 (C12, red) overnight with or without atglistatin (ATGLi) or lalistat 2 (Lali2) followed by staining with Bodipy 493 (green) and DAPI (white). (B) Number and (C) size of Bodipy C12 positive lipid depots. Data from three independent experiments are combined taking the average value of DMSO treated Cont cells as 1. n=18 to 26 cells total in (B), yielding n= 73 to 506 lipid depots in (C). Mean is indicated. Y axis is log 10 for (B) due to wide distribution of data. (D) TG contents corrected for protein contents and expressed taking value for DMSO treated Cont as 1. Mean ± sem, n=4-8. (E, F) Cont and PLIN2KD INS-1 cells metabolically labeled with C12 overnight followed by immunostaining of mitochondria by anti-HSP60 antibody. A part of PLIN2KD cells were incubated with Lali2 overnight. (E) Pearson’s coefficient for C12 and HSP60 was calculated as in methods. n=15. Representative data from four independent experiments. Mean ± sem. (F) Representative confocal image. *; *p*<0.05 by One-way ANOVA test with posttest by Sidak’s test for B, D, Krusaki-Wallis test for C, and Dunnett’s test for E.

### The inhibition of lysosomal acid lipase prevents mitochondrial fragmentation in PLIN2 KD INS-1 cells and human beta cells

We next addressed whether Lali2 prevents the fragmentation of mitochondria in PLIN2 KD beta cells. Form factor, aspect ratio, and mitochondria length were all decreased in PLIN2KD INS-1 cells compared with Cont cells indicating that mitochondria are fragmented. Lali2 treatment effectively reversed all three parameters significantly in PLIN2KD INS-1 cells (Fig. 5A-D). Similar prevention of mitochondrial fragmentation was seen in PLIN2 KD human beta cells treated with Lali2 (Fig. 5E). Form factor and mitochondria length were reduced in PLIN2 KD beta cells and increased in Lali2 treated PLIN2 KD beta cells (Fig. 5F, G). Thus, lysosomal degradation of LD is required for both increased flux of Bodipy C12 to mitochondria and mitochondrial fragmentation in PLIN2 KD beta cells.

**Figure 5.**
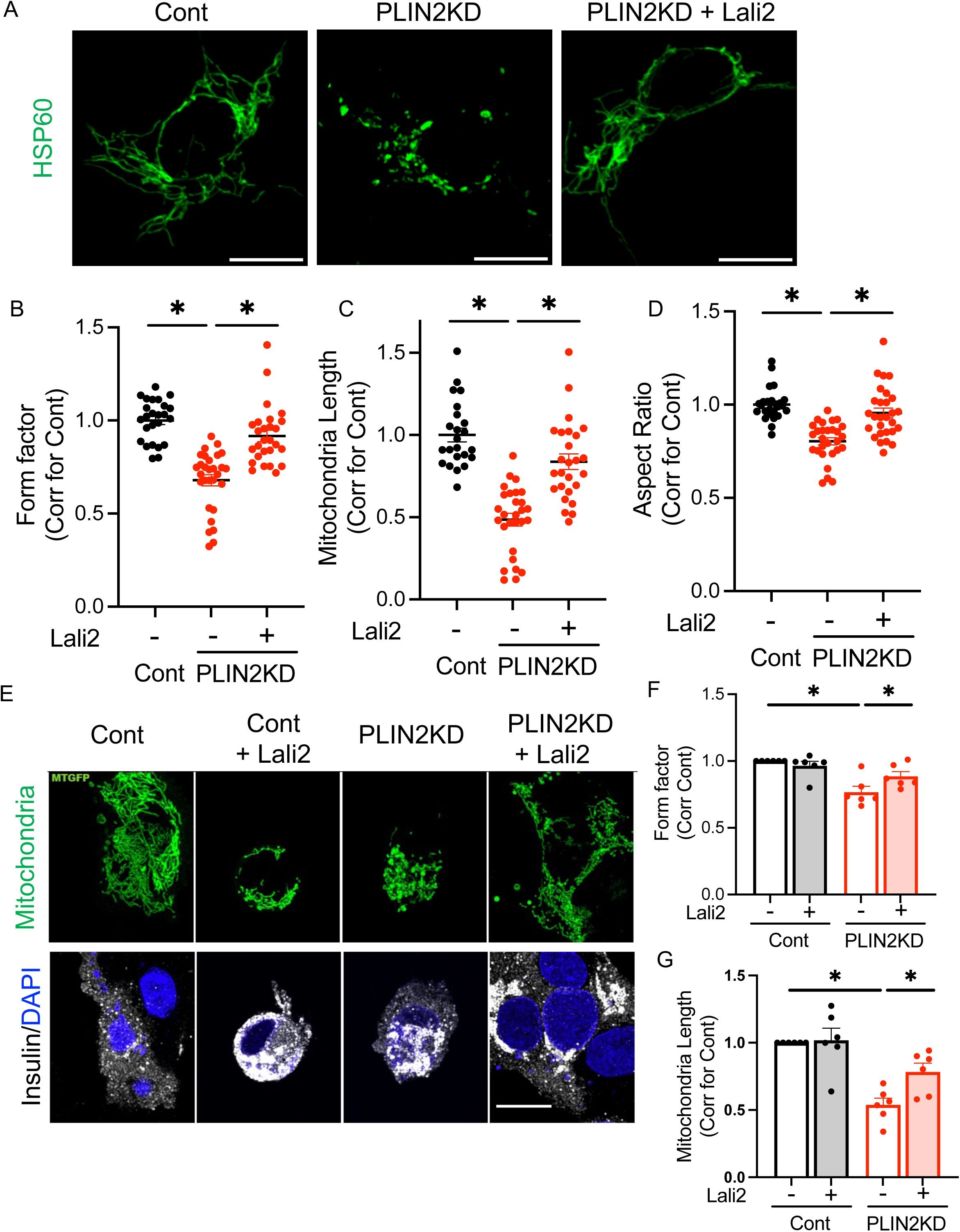
A LIPA inhibitor prevents fragmentation of mitochondria in PLIN2 downregulated INS-1 cells and human beta cells. (A-D) Mitochondria were visualized by anti-HSP60 antibody in control (Cont) and PLIN2 deficient (PLIN2KD) INS-1 cells. A part of PLIN2KD cells were treated with lalistat 2 (Lali2) overnight prior to fixation. (B) Form factor, (C) mitochondrial length, and (D) aspect ratio were obtained as parameters of mitochondrial morphology as in methods. Data combine three independent experiments and expressed by taking the average of Cont as 1 for each experiment. Each dot represents one cell and n= 23 to 29 cells were counted in total. (E-F) Mitochondria (mitochondria-GFP), beta cells (anti-insulin antibody, white) and nuclei (DAPI, blue) were visualized in human islet cells treated with control (Cont) and lenti-shPLIN2 (PLIN2KD) with or without Lali2. (E) Representative images. (F) Form factor and (G) mitochondrial length obtained as in methods in five donor islets. Data were expressed by taking the average value of DMSO treated Cont as 1 for each donor and each dot represents one donor. All scale bars are 10 μm. Data represent mean ± SEM *; *p*<0.05 by one-way ANOVA. Tukey’s posttest for (B-D). Sidak posttest for (E and F).

### Lipid droplet-lysosome interaction is increased in PLIN2KD INS-1 cells

While macroautophagy has been shown to degrade LDs,^33^ recent studies have shown that lysosomes can directly contact LDs to digest LD contents in hepatocytes.^34^ Split-GFP-based contact site sensor (SPLICS) allows the detection of an organelle contact by targeting two complementary parts of green fluorescent protein (GFP) to the surface of two organelles of interest.^35^ We tagged livedrop that distributes to LD surface^36^ with GFP_1-10_ (validation in supplementary Fig. 2A) and TMEM192 that distributes to lysosome surface^37^with β_11_ (validation in supplementary Fig. 2B) to form a full GFP when a LD and a lysosome are in the close vicinity (SPLICS-P2A-LD–LYSO, Fig. 6A). When SPLICS-P2A-LD–LYSO was expressed in PLIN2 KD INS-1 cells and Cont cells, PLIN2 KD INS-1 cells had more green dots indicating the increase in LD-lysosome contacts (Fig. 6B, C). PLIN2 KD in INS-1 cell and human pseudoislets did not upregulate the expression of transcription factor EB (TFEB), a key transcription factor that regulates the autophagy-lysosome pathway (Fig. 6D, E).^38^ However, the expression of *LIPA* was upregulated in both INS-1 cells and human pseudoislets without a change in *PNPLA2* expression indicating a selective impact of PLIN2 KD on *LIPA* expression (Fig. 6D, E). 3-methyladenine (3MA) that inhibits Class III PI-3 kinase, a kinase with a role in autophagosome formation,^39^ did not prevent the increase of Bodipy C12 in mitochondria (Fig. 6F) or the fragmentation of mitochondria based on form factor (Fig. 6G), mitochondrial length (Fig. 6H), and aspect ratio (Fig. 6I) in PLIN2 KD INS-1 cells. Thus, autophagosome formation is not required for the delivery of Bodipy C12 to mitochondria or the fragmentation of mitochondria in PLIN2 KD cells. PLIN2 KD was not sufficient to increase overall autophagy flux either considering that LC3-II/I level was similar in the presence and absence of chloroquine (CQ) between Cont and PLIN2 KD cells (Fig. 6J, K). The reduced activity of mTOR is a strong stimulus of macroautophagy.^40^ However, phosphorylation of ribosomal protein S6, a marker of mTORC1 activity was not altered in PLIN2 KD INS-1 cells (Supplementary Fig. 2C, D). Collectively, data do not support the involvement of macrolipophagy. The increase in direct interaction between LD and lysosome strongly indicates that microlipophagy is upregulated by PLIN2 KD and mediates the transfer of lipids from LD to lysosome in PLIN2 KD beat cells.

**Figure 6.**
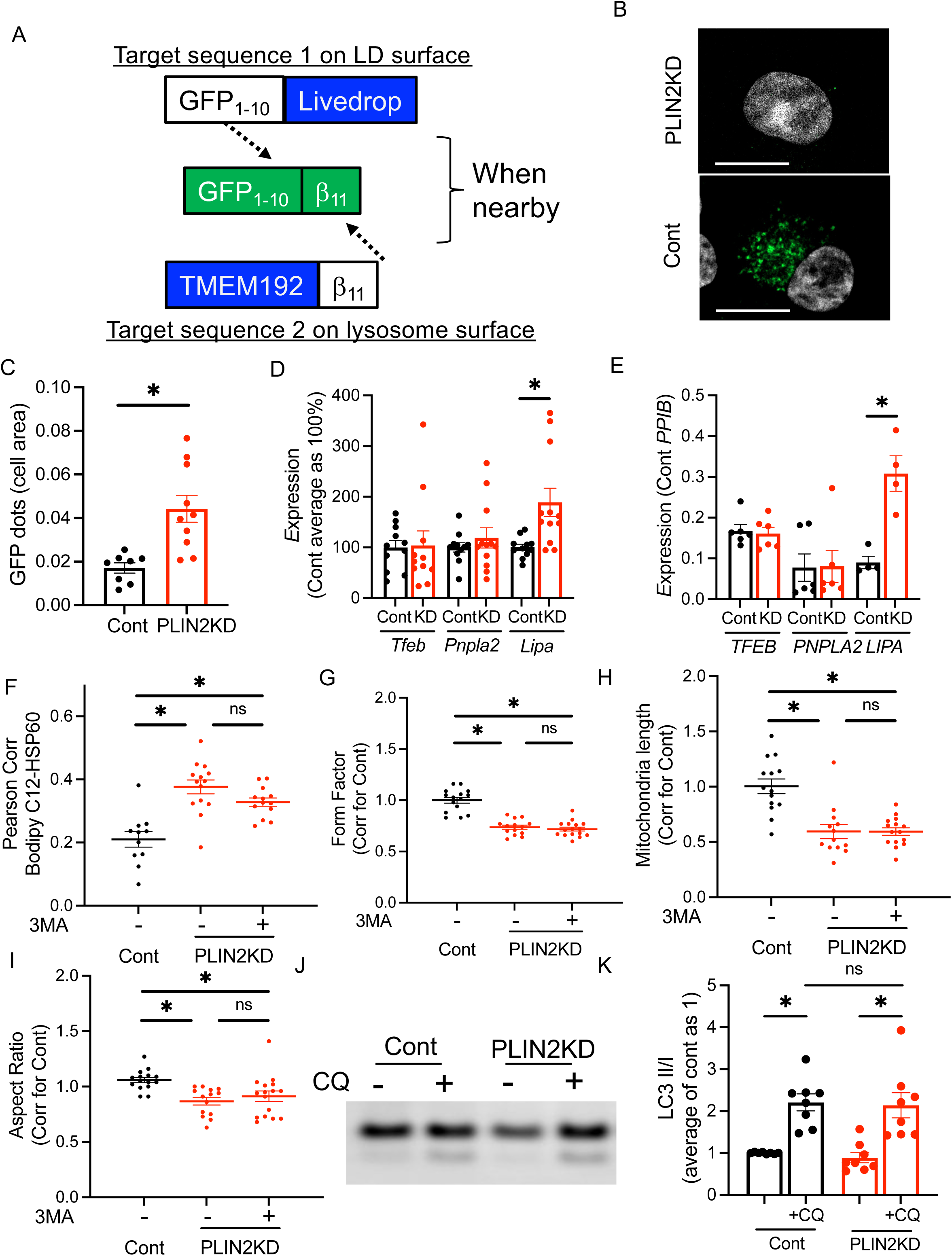
Direct interaction between lysosome and lipid droplets is increased in PLIN2 deficient INS-1 cells. (A) Scheme of a lipid droplet (LD)-lysosome split-GFP-based contact site sensor (SPLICS-P2A-LD–LYSO). Livedrop tagged with GFP_1-10_ is expressed on the surface of LD and TMEM192 tagged with β_11_ is expressed on the surface of lysosome. When they are within 10 nm vicinity, GFP_1-10_ and β_11_ fuse to form a GFP. (B, C) Number of green dots corrected for cell area was compared between control (Cont) and PLIN2 downregulated (PLIN2KD) INS-1 cells that were transduced by lentivirus expressing SPLICS-P2A-LD–LYSO. n= 8-10 images. (B) Representative image and (C) representative data from four independent experiments. (D, E) RT-qPCR determined the expression of genes indicated in (D) INS-1 cells (n=11 to 12) and (E) human pseudoislets (n=4 to 6 donors) with or without downregulation (KD) of PLIN2. (F-I) Cont and PLIN2KD INS-1 cells were treated with or without 3.5 mM 3MA overnight in the presence of Bodipy C12. Mitochondria were labeled with anti-HSP60 antibody. (F) Pearson’s coefficient for Bodipy C12 and HSP60. Each dot represents one cell; n=11 to 14 cells combined from two independent experiments. (G) Form factor, (H) mitochondria length, and (I) aspect ratio in Cont and PLIN2KD INS-1 cells expressed taking the average of Cont as 1. Each dot represents one cell and n=13 to 16 cells from two independent experiments are combined. (J, K) Western blot comparing LC3I and LC3II abundance in Cont and PLIN2KD INS-1 cells treated with or without chloroquine (CQ). (J) Representative blot and (K) the ratio of LC3II/I from densitometry obtained from three independent experiments and is expressed taking the average of Cont without CQ as 1. n=8. Data are mean ± SEM. *; *p*<0.05 by student’s t-test for (C-E, K), and One-way ANOVA for the rest.

### Lysosome-mitochondria interaction is increased in PLIN2 KD INS-1 cells

In hepatocytes, serum starvation triggers lipophagy, increases beta oxidation of FA (FAO), and accumulates BodipyC12 in mitochondria.^41^ Interestingly, the increased trafficking of FA to mitochondria in hepatocytes under starvation was preceded by the efflux of FA through lysosomal secretion (Fig. 7A left).^41^ With the similar localization of Bodipy C12 in mitochondria and upregulation of lipophagy in PLIN2KD INS-1 cells, we asked whether Bodipy C12 secretion is required for PLIN2 KD INS-1 cells to increase Bodipy C12 in mitochondria and the fragmentation of mitochondria. In contrast to starved hepatocytes, Bodipy C12 secretion was not increased in PLIN2 KD INS-1 cells (Fig. 7B). Also, trapping secreted FA in medium by FA-free BSA or blocking re-entry of FA by CB16.2 (Fig. 7A) did not prevent the increase of Bodipy C12 in mitochondria (Fig. 7C) or mitochondrial fragmentation (Fig. 7D). Rather, the contacts between lysosome and mitochondria were increased in PLIN2 KD INS-1 cells. There was no change in mitochondrial mass or lysosome mass (Supplementary Fig. 3A, B). Thus, the close contact between lysosome and mitochondria likely contributes to Bodipy C12 accumulation and mitochondrial fragmentation in PLIN2 KD INS-1 cells (Fig. 7A right). Importantly, blocking lysosomal degradation of TG was sufficient to improve insulin secretion in PLIN2 KD INS-1 cells without change in insulin contents (Fig. 7G, Supplementary Fig. 3C).

**Figure 7.**
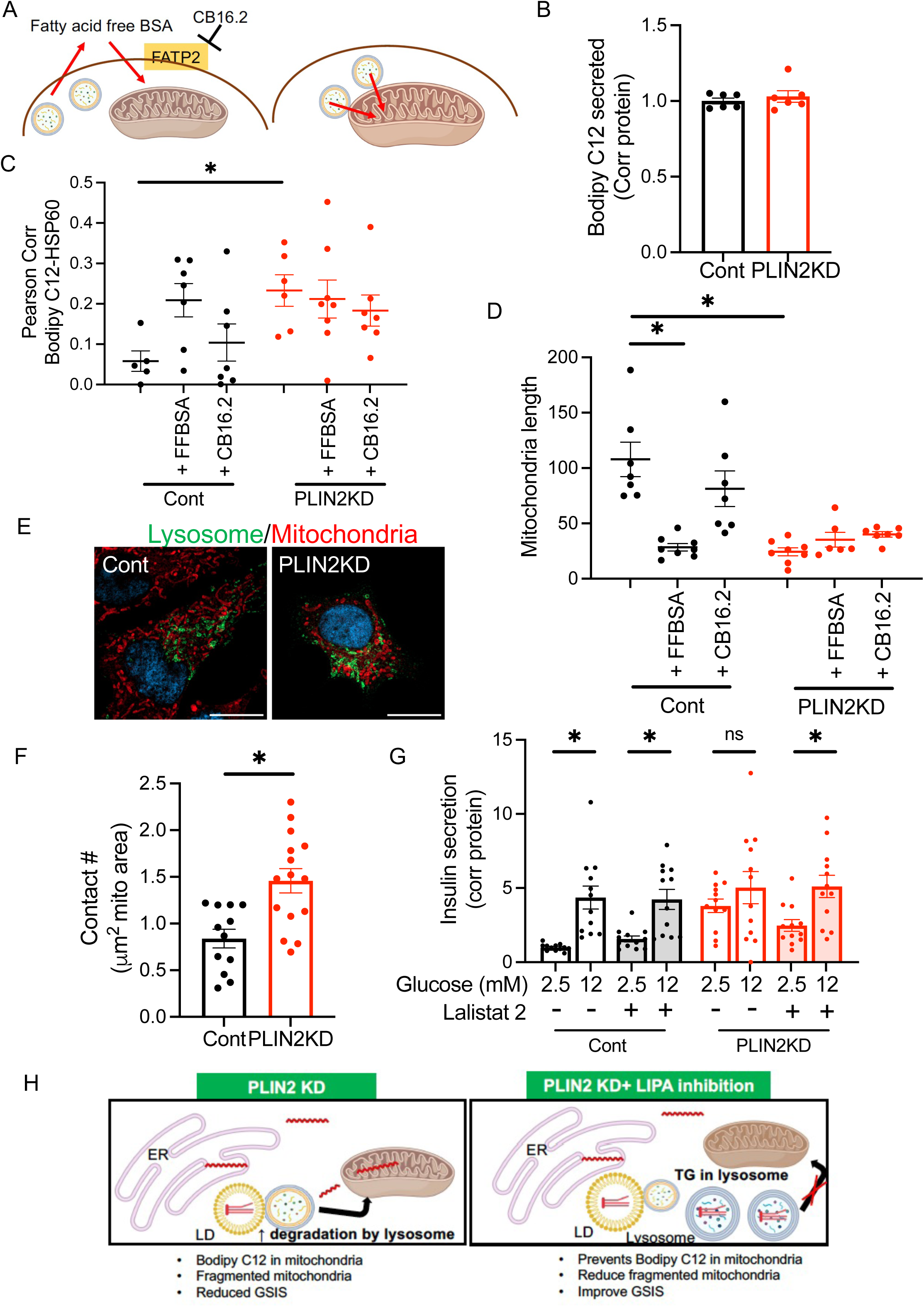
Lysosome-mitochondria contact is increased in PLIN2 deficient INS-1 cells. (A) Two potential pathways by which FA are transferred from lysosomes to mitochondria. On the left, FA are secreted to the medium by the lysosome and taken up by the cells prior to entering mitochondria as shown in serum-starved hepatocytes.^41^ This step was shown to be inhibited by either binding released FA to fatty acid-free BSA (FFBSA) or by blocking FA reentry using FTAP2 inhibitor CB16.2.^41^ On the right, FA are directly transferred from lysosome to mitochondria through contacts. (B-D) Control (Cont) and PLIN2 downregulated (PLIN2KD) INS-1 cells were labeled with Bodipy C12 (C12) overnight and chased for 4 h to measure the secretion of C12 as described in methods. Cont and PLIN2KD INS-1 cells were fixed and immunostained for mitochondria by anti-HSP antibody. (B) C12 in the medium at the end of 4 h was corrected for protein contents in cell lysate and expressed taking the average of Cont as 1. n=6 combined from three independent experiments each in duplicates. (C) Pearson’s coefficient for C12 and HSP60 and (D) mitochondria length in Cont and PLIN2KD cells treated with or without FFBSA or CB16.2. n=5 to 8 images for (C) and 6 to 8 images for (D), representative data of two independent experiments. (E) Representative images and (F) contact number between lysosome marked by CellLight Lysosomes-GFP (green) and mitochondria marked by MitoTracker deep red in Cont and PLIN2KD INS-1 cells. n= 12 to 15 images. Data is representative of three independent experiments. (G) Cont and PLIN2 KD INS-1 cells were cultured with or without lalistat 2 (Lali2) for 16 h and insulin secretion was measured by static incubation. Data were corrected for protein contents and expressed taking the average of Cont at 2.5 mM glucose as 1. n=12 combining four independent experiments. (H) Diagram showing FA trafficking between LD, lysosome and mitochondria in PLIN2 KD beta cells with or without Lali2. Data represent mean ± SEM. *; *p*<0.05 by one-way ANOVA for (C, D, G) or Student’s t-test for (F). ns: not significant.

### LIPA inhibition prevents mitochondrial changes and improves glucose-stimulated insulin secretion in beta cells exposed to the glucolipotoxic condition

The exposure to elevated glucose and FA is known to increase LD formation and PLIN2 levels in beta cell models.^16,18,19^ However, the expansion of LDs may cause relative insufficiency of PLIN2 and may increase the access of lysosomes to LDs for lipophagy. We compared TG contents in the presence and absence of Lali2 to determine the proportion of TG that is degraded by Lali2 in INS-1 cells cultured in regular medium and the medium with higher levels of glucose and FA (GLT). Lali2 increased TG in INS-1 cells significantly under GLT condition indicating that LIPA actively degrades a substantial portion of TG under GLT condition (Fig. 8A). Also, two hallmarks of PLIN2-KD beta cells, the increase in Bodipy C12 in mitochondria and the mitochondria fragmentation were also seen in INS-1 cell cultured under GLT condition (Fig. 8B-F, Supplementary Fig. 4A). Moreover, Lali2 was sufficient to prevent these changes indicating that the inhibition of lysosomal degradation of LDs prevents mitochondria changes in INS-1 cells under GLT condition as well (Fig. 8B-F, Supplementary Fig. 4A). Lali2 was also effective in preventing mitochondrial fragmentation under GLT in human beta cells (Fig. 8G-I, Supplementary Fig. 4B). Lastly, we tested whether Lali2 improves insulin secretion in human islets cultured under GLT condition overnight. Compared with control culture condition, GLT condition reduced insulin contents (Cont without lali2 vs. GLT without lali2 in Fig. 8J), increased basal insulin secretion (Cont without lali2 vs. GLT without lali2 in Fig. 8K), and markedly reduced GSIS (Cont without lali2 vs. GLT without lali2 in Fig. 8L), changes well recognized to occur in beta cells under nutrient load.^42–45^ While LIPA inhibition in mouse islets is reported to increase in GSIS,^30^ the inhibition of LIPA in human islets did not increase GSIS in control islets (Cont without lali2 vs. Cont with lali2 in Fig. 8L). Lali2 did not significantly improve changes in insulin contents or basal insulin secretion in human islets under GLT condition (GLT without lali2 vs. GLT with lali2, Fig. 8 J, K). However, Lali2 significantly improved glucose responsiveness of insulin secretion (GLT without lali2 vs. GLT with lali2, Fig. 8L), which is also demonstrated as the improvement of stimulation index (Fig. 8M). Thus, the blocking lysosomal degradation of LDs prevents fragmentation of mitochondria and improves GSIS in human beta cells exposed to GLT.

**Figure 8.**
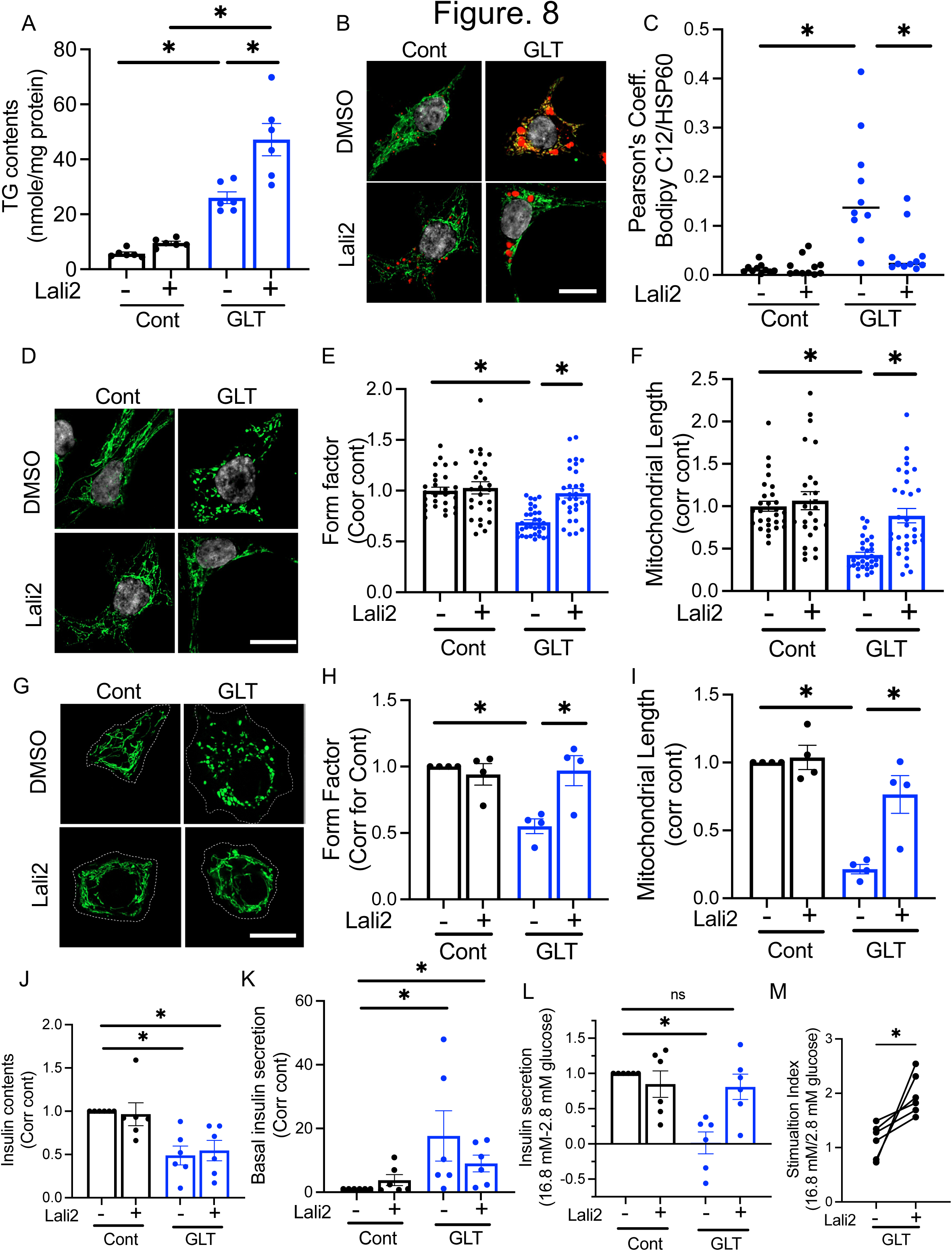
A LIPA inhibitor prevents mitochondria fragmentation and improves glucose-stimulated insulin secretion in beta cells under glucolipotoxic condition. (A) INS-1 cells were cultured in RPMI medium containing 11.1 mM glucose (Cont) or 20 mM glucose + 0.16 mM oleic acid (OA) + 0.32 mM palmitic acid (PA)(GLT) in the presence or absence of 20 μM lalistat 2 (Lali2) for 48 h. Triglycerides (TG) contents at the end of incubation was measured and corrected for protein contents. n=6, combining duplicates from three independent experiments. (B) Representative images of Bodipy C12 (C12) and HSP60, (C) Pearson’s coefficient for C12 and HSP60, (D) representative image of mitochondria, (E) form factor, and (F) mitochondria length of Cont and GLT INS-1 cells treated for overnight with or without Lali2 in the presence of C12. After fixation, mitochondria were visualized by anti-HSP antibody. n= 28 to 32 images from three independent experiments. (G) Representative image, (H) form factor and (I) mitochondria length of human beta cells cultured in CMRL1066 containing 5.5 mM glucose (Cont) or CMRL1066 at 20 mM glucose + 0.16 mM OA+ 0.32 mM PA (GLT) with or without 10 μM Lali2 overnight. Mitochondria were visualized by CellLight Mitochondria-GFP. Beta cell border identified by anti-insulin antibody is indicated by dotted line. Each dot in figures represents the average of values from each donor, n= 4 donors. (J-M) Human islets were cultured overnight in CMRL1066 medium containing 5.5 mM glucose (Cont) or 12.5 mM glucose + 0.8 mM OA+ 0.16 mM PA (GLT) with or without 10 μM Lali2 overnight. Insulin secretion at 2.8 mM glucose (basal) and 16.8 mM glucose was measured as in methods. (J) Total insulin contents per IEQ, (K) basal insulin secretion, (L) the increment of insulin secretion at 16.8 mM glucose, and (M) stimulation index were expressed taking Cont without Lali2 for each donor as 1. n=6 donors. Data represent mean ± SEM *; *p*<0.05 by one-way ANOVA (A-L) or Student’s t-test (M). n.s: not significant

## DISUCSSION

KO or KD of PLIN2 impairs GSIS in multiple models of beta cells including INS-1 cells, EndoC cells, mouse beta cells, and human islets indicating that PLIN2 is indispensable for the proper function of beta cells.^18,19^ The previous study showed mitochondrial parameters are altered in PLIN2 deficient beta cells: the reduction of OXPHOS proteins (INS-1 cells, mouse islets, human islets), low OCR (INS-1 cells, mouse islets), the increase in mid chain acylcarnitines (INS-1 cells), fragmented mitochondria (INS-1 cells), and Bodipy C12 accumulation in mitochondria (INS-1 cells) were seen after PLIN2 downregulation.^18^ Here, we demonstrated that the fragmentation of mitochondria was also seen in human beta cells but not in alpha cells after PLIN2 KD. PLIN2 KD in human islets impaired insulin secretion while the regulation of glucagon secretion was grossly preserved. Data collectively support that mitochondria are a target by which PLIN2 deficiency impairs insulin secretion. In the current study, the inhibition of lysosomal degradation of lipids was sufficient to prevent Bodipy C12 localization in mitochondria, mitochondrial fragmentation, and the impaired insulin secretion in beta cell models indicating that the accelerated lipophagy impairs insulin secretion through FA delivery to mitochondria in PLIN2 KD beta cells (Fig. 7H). Both LD-lysosome and lysosome-mitochondria contacts were increased in PLIN2 KD INS-1 cells implicates that FAs are relayed through three organelles and PLIN2 regulates this FA flow in beta cells. More importantly, we found that LIPA inhibition prevents mitochondrial fragmentation and improves GSIS in human beta cells exposed to GLT condition revealing a new role of lipophagy in beta cell demise under nutrient overload.

PLIN family of proteins including PLIN2 is considered to alter the size and number of LD predominantly by regulating LD degradation rather than synthesis of LDs. However the precise mechanism utilized by PLIN to regulate LD degradation varies depending on the type of cells, conditions of cells, and the type of PLIN being expressed.^15^ PLIN1 and PLIN5 are well known to regulate lipolysis by modulating access of PNPLA2 and its co-factor ABHD5 to LDs.^15^ Although PLIN2 does not interact with PNPLA2 or ABHD5 directly, it is considered to reduce access of PNPLA2 to LDs. In fibroblasts and hepatocytes under serum starvation, access of PNPLA2 to LDs was increased after removal of PLIN2 by chaperon mediated autophagy.^46^ PLIN2 is also proposed to regulate macroautophagy of LDs in hepatocytes.^47,48^ In the current study of beta cells under the nutrient sufficient condition, the inhibition of PNPLA2 partially restored the size of LDs in PLIN2 KD INS-1 cells indicating that PLIN2 in beta cells prevents degradation of LDs by both PNPLA2 and lipophagy. However, it was the inhibition of LIPA that prevented Bodipy C12 accumulation in mitochondria highlighting a specific role lipophagy in directing FA to mitochondria in beta cells.

The increased lipophagy and the localization of Bodipy FA probe in mitochondria were previously reported in serum starved hepatocytes/fibroblasts and glioblastoma.^41,49^ These are cells with high demand for FAO and lipophagy is considered to supply FA to mitochondria to support FAO in this case.^41,49^ PLIN2 level is posttranslational regulated by proteasomal degradation of PLIN2 that is not bound to LDs.^50^ Thus, PLIN2 level will decline when FA availability for TG synthesis is reduced. This provides a mechanism to regulate lipophagy depending on beta cell’s energy state. Under a glucose deprived condition in beta cells, the reduction of PLIN2 is expected to allow the utilization of neutral lipids stored in LDs for FAO and aids the adaptation of cells to the glucose deprived condition. However, genetic PLIN2 KO/KD in beta cells creates a condition where the access of lysosomes to LDs is increased under a nutrient sufficient condition dissociating the demand for FAO from the influx of FA to mitochondria via lipophagy. Limiting FA delivery to mitochondria under a glucose sufficient condition is likely very important for beta cells considering that beta cells favor glucose oxidation over FAO to support GSIS.^51^ While carnitine palmitoyl transferase 1a (CPT1a) plays an important role in sensing glucose levels through malonyl-CoA and suppresses FA entry to mitochondria in beta cells, the suppression of CPT1a by malonyl-CoA is not complete and can be reduced by exposure to FA.^52,53^ Thus, PLIN2 may provide an additional layer of protection to beta cells to prevent excessive influx of FA to mitochondria through lipophagy. Interestingly, the FA flow from LD to mitochondria through lipophagy appears to contribute to mitochondria fragmentation and impaired GSIS under GLT condition in human beta cells without PLIN2 KD. It appears that PLIN2 does not control lipophagy sufficiently when LD pool expands despite the increase in PLIN2 levels in beta cells under nutrient loading.

GLT condition elicits broad effects on beta cells through multiple pathways. Thus, it is not surprising that the inhibition of LIPA improved GSIS in human islets cultured under GLT condition but did not prevent the reduction of insulin contents or the increase in basal insulin secretion in the current study. The reduction of insulin contents and the increase of basal insulin secretion may not be due the delivery of FA to mitochondria by lipophagy. Although a mechanism is yet to be determined, the reduction of INS contents we observed is likely a part of the programmatic reduction of translation of genes important for insulin secretion reported in beta cells and pancreatic islets chronically exposed to glucose.^43^ Multiple mechanisms are proposed for the elevated basal insulin secretion under nutrient load in beta cells including the increase in reactive oxygen species (ROS) and cytosolic long chain-CoA, S-palmitoylation of K_ATP_ channel, and protein leak in mitochondria.^44,45^ In contrast, mitochondrial metabolism of glucose is considered to be one of the major determinants of glucose responsiveness of insulin secretion.^54^ The current study indicates that the excessive lipophagy under nutrient load may impair glucose responsiveness of beta cells by exposing mitochondria to FA.

With the potential relevance of lipophagy in beta cell dysfunction under nutrient load, it is of interest to understand how FA are transferred between LD-lysosome and lysosome-mitochondria in beta cells both after PLIN2 KD and under GLT condition. The current study showed that the direct contact between LDs and lysosomes is increased in PLIN2 KD INS-1 cells indicating that microlipophagy mediates transfer of LD contents to lysosome without the formation of autophagosome. In hepatocytes, microlipophagy has been shown to target small LDs.^55^ Considering a relatively small size of LDs in beta cells, the encasement by autophagosome may not be needed for beta cell lipophagy. Lysosome-mitochondria interaction in our model did not involve lysosomal secretion that mediates FA transfer to mitochondria in serum starved hepatocytes/fibroblasts.^41^ Direct transfer of FA from lysosome to mitochondria may be needed for efficient FA transfer in beta cells due to a significantly smaller pool size of LDs compared with hepatocytes. While the current study strongly supports that the direct LD-lysosome-mitochondria contact mediates transfer of FA, an additional support can be obtained if the disruption of intact prevents FA movement between lysosomes and mitochondria. For that, further studies are needed to clarify a molecular tether that mediates the direct interaction between LD-lysosome and lysosome-mitochondria in beta cells.

It is important to note that PLIN2 could play different roles depending on cells and the condition of cells. Similar to beta cells, the downregulation of PLIN2 in human dermal fibroblasts impaired mitochondrial function.^56^ In contrast, hepatocyte specific KO of PLIN2 improves metabolic phenotypes of diet-induced steatosis and steatohepatitis in mice.^48,57^ In addition to regulating lipid trafficking, PLIN2 is considered to organize defensive proteins on LDs under infection.^58^ Thus, it is importance to address cell type specific roles of PLN2.

The current study has several limitations that benefit future studies. Although cultured beta cells allow detail analyses of Bodipy C12 trafficking, mitochondrial morphometry, and organelle interaction, the contribution of lipophagy in beta cell dysfunction in PLIN2 KD beta cells and under GLT condition should be tested in vivo model. As LD morphology is significantly different in mouse beta cells, mice may not serve as an ideal model for in vivo study.^32^ LIPA mediates hydrolysis of cholesterol ester as well.^28^ Although the production of cholesterol ester in beta cells is low (data not shown), the further study is required to determine whether Lali2 alters cholesterol homeostasis in beta cells. The increase in GCG contents is a novel phenotype noted by PLIN2 KD in human islets and warrants further study to address a mechanism and its impact on the regulation of glucagon secretion beyond a limited study we performed here.

In summary, PLIN2 in beta cells regulates FA trafficking to mitochondria through lipophagy and is important for preserving GSIS. FA trafficking from LD to mitochondria through lipophagy also contributes to impaired GSIS under nutrient load in beta cells and serves as a potential target to improve glucose responsiveness of beta cells under nutrient load.

## METHODS

### Processing of human islets

Institutional Review Board at University of Iowa deemed human islet experiments are not a human study. Human islets from Integrated Islet Distribution Program (IIDP), Alberta Diabetes Institutes or Prodo Laboratories (Alliso Viejo, CA) with reported viability and purity above 80% were cultured in CMRL1066 containing 1% human serum albumin, 1% Pen-Strep, and 1% L-glutamate overnight at 37 °C and 5% CO_2_ upon arrival for recovery from shipping. The creation of human pseudoislets treated with lentivirus expressing shRNA against PLIN2 and scramble control were previously reported.^18^ Downregulation of PLIN2 was monitored by rt-PCR. For GLT condition, indicated concentration of oleic acids (OA) and palmitic acids (PA) mixture at 1:2 ratio was added to medium containing indicated concentration of glucose. Treatment of human islets/islet cells with Lali2 was at 10 μM overnight unless specified otherwise.

### INS-1 cell culture

823/13 cells (INS-1 cells, a kind gift from Dr. Christopher Newgard (Duke University)) were maintained in RPMI1640 with 10 mM HEPES, 2 mM L-glutamine, 1 mM sodium pyruvate, 50 μM β-mercaptoethanol, and penicillin+ streptomycin (INS-1 medium) supplemented with 10% FBS. For down-regulation of PLIN2, cells were transfected with 30 nM of DsiRNA targeting PLIN2 (rn.Ri.Plin2.13.1 from Integrated DNA Technologies, Coralville IA) using DharmaFECT1 transfection reagent (Horizon Discovery, Lafayette, CO) as previously validated and reported.^18^ Non-targeting DsiRNA (Integrated DNA Technologies) was used as negative control. Alternatively, PLIN2 was downregulated by lentivirus carrying shRNA sequence targeting rat PLIN2 (CAGAAGCTAGAGCCGCAAATT). Lentivirus expressing scramble (CCTAAGGTTAAGTCGCCCTCG) RNA was used as control. Downregulation of PLIN2 was monitored by rt-PCR. For Western blot of LC3A/B, a part of cells was treated with 50 μM CQ for 2 h prior to harvest. For GLT condition, indicated concentration of OA and PA mixture at 1:2 ration was added to INS-1medium containing indicated concentration of glucose. Treatment of INS-1 cells with Lali2 and Atglistatin was both at 20 μM overnight unless specified otherwise.

### mRNA and quantitative RT-PCR

RNA was isolated from INS-1 cells using RNeasy kit (Qiagen, Germantown, MD) and from islets using TRIzol reagent (ThermoFisher Scientific, Waltham, MA) and cDNA was synthesized as published.^29^ Gene expression was assessed as previously described using ABI TaqMan commercial primers (ThermoFisher Scientific).^29^

### Perifusion of Islets

BioRep Perifusion System (BioRep Technologies, Miami Lakes, FL) was used to perifuse human islets and pseudoislets as published.^59^ For the measurement of insulin secretion, islets were perifused by Krebs-Ringer bicarbonate buffer (KRB) containing 2.8 mM glucose for 48 minutes followed by KRB containing 16.7 mM glucose or 30 mM KCl plus 2.8 mM glucose for indicated time. For the measurement of glucagon secretion, islets were perifused by KRB containing 3.3 mM glucose for 48 min followed by 1 mM glucose plus amino acid mixture (2 mM each of glutamine, alanine, and arginine as published in^60^) for 28 min and 7 mM glucose plus amino acid mixture for 14 min. Total insulin and glucagon contents were obtained from islets incubated overnight at 4 °C in RIPA buffer (R0278, Millipore Sigma, St Louis, MO) containing protease inhibitors. Insulin was measured using STELLUX Chemiluminescent Human Insulin ELISA (ALPCO, Macedon NY). Glucagon was measured using glucagon ELISA kit from Crystal Chem (Elk Grove Village, IL). Hormone secretion at the specified concentration of glucose and KCl was measured as area under curve (AUC) of each peak divided by the length of peak (min).

### Preparation of human islet cells and INS-1 cells for image analyses

Human islets dispersed by Accutase (SCR005, Millipore Sigma, St Louis, MO) were plated at approximately 35,000 cells/cm^2^ on a confocal dish coated with 50 μg/mL of collagen IV (C5533 from Sigma) as reported,^23^ and cultured in neuronal medium (11 mM glucose) described by Phelps et al.^22^ at 37 °C at 5% CO_2_ for three days prior to fixation. INS-1 cells were grown on a cover glass coated with extracellular matrix of HTB9 cells and cultured in INS-1 cell medium at 37 °C at 5% CO_2_ for overnight to three days prior to fixation. Both human islet cells and INS-1 cells were fixed in 4% paraformaldehyde as published.^18^

For both human islets and INS-1 cells, 2 to 3 μM of Bodipy C12 was added during culture to monitor the spatial distribution of this Bodipy C12 (30 min to 6 h) and to visualize TG rich LDs (overnight).^61^ To visualize mitochondria in human beta cells, human islet cells cultured on a confocal dish were transduced with CellLight Mitochondria-GFP, BacMam 2.0 (ThermoFisher) at 50 particle per cell (PPC) two days before harvest.

### Pearson’s coefficient and morphometry of LDs and mitochondria

Human islet cells were immunostained for INS (rabbit anti-INS antibody, C27C9, Cell signaling at 1:300) or GCG (mouse anti-GCG antibody, K79bB10 from Sigma at 1:600) to visualize beta and alpha cells, respectively using a published protocol.^23^ Mitochondria in INS-1 cells were visualized using mouse anti-HSP60 antibody (66-41-1-lg, Protein Tech, Rosemont, IL) at 1:300. To visualized neutral lipids, fixed cells were incubated with 2 μM Bodipy 493/503 (Bodipy 493, ThermoFisher) as published.^62^ Nuclei was visualized by 1 μg/ml 4′,6-diamidino-2-phenylindole (DAPI, from ThermoFisher) as published.^23^

Zeiss 980 microscope Airy Scan 2 in an Airy scan SR mode was utilized to acquire z-stack images via a 63x oil lens at an interval of 0.15 µm covering the entire height of INS-1 cells and human islet cells. For human islet cells that are more cuboidal than INS-1 cells, 2 sets of 5 consecutive slices of Z-stack image were selected to represent bottom half and top half of each islet cluster. 5 to 10 islet clusters were analyzed for each condition. Thereafter, 5 consecutive slices were merged into one image via the maximum-intensity projection plugin in ImageJ/Fiji. Size and number of LDs defined by Bodipy 493 or Bodipy C12 were determined using Image J based program as published.^23^ Pearson’s coefficient of Bodipy C12 and mitochondria were calculated using the colocalization function in Bitplane Imaris v 9.6 (Oxford Instruments, Concord, MA). Mitochondrial images captured by a Zeiss LSM980 microscope were converted to binary images of mitochondrial particles using a custom written NIH ImageJ Marco as published ^63,64^. Automated morphometry of mitochondrial particles was further performed using the NIH ImageJ Marco to obtain aspect ratio (major axis/minor axis), form factor (perimeter^2^/(4π x area)), and mitochondria length.^63,64^ For human islet cells, area of beta cells (INS^+^) and alpha cells (GCG^+^) was also obtained by ImageJ/Fiji to normalize data.

### TG contents

INS-1 cells were solubilized in RIPA buffer with protease inhibitors as published.^29^ Protein contents of cell lysate was measured by Pierce BCA protein assay (ThermoFisher). Then, TG was extracted from cell or islet lysate by Folch buffer as published^16^ and quantitated by Infinity Reagents (ThermoFisher). TG contents were corrected for protein contents.

### Static incubation

After 1 h pre-incubation in glucose free KRB, INS-1 cells were incubated for 1 h in KRB containing the indicated concentration of glucose. Insulin secreted and insulin contents were measured with STELLUX Chemiluminescent rodent Insulin ELISA respectively (ALPCO). Lysates of INS-1 cells in RIPA buffer was prepared as for TG contents above.

### Expression of SPLICS-P2A-LD–LYSO and measurement of LD-lysosome contacts

To detect contacts between LDs and lysosome, split-GFP-based contact site sensors (SPLICS) was designed by modifying ER-MITO-SPLICS published.^35^ LD targeted GFP_1-10_ and lysosome targeted β_11_were connected by a 2A peptide for equimolar expression of both constructs. For LD targeting, livedrop was amplified from pKeima-LiveDrop, a kins gift from Drs. Tobias Walther and Robert Farese (Addgene plasmid # 199695).^36^ TMEM192 was amplified from pMRX-IB-TEME192 mGFP, a kins gift from Dr. Noboru Mizushima (Addgene plasmid # 184897).^65^ Lentivirus was made by Vectorbuilder (Chicago, IL). INS-1 cells treated with siRNA targeting PLIN2 or non-targeting siRNA were plated on a cover glass and transduced by lenti-SPLICS-P2A-LD–LYSO. After fixation as above, confocal image of cells was capture by Zeiss 980 microscope and number of green dots per cell area was obtained.

### Western Blot

Lysates of INS-1 cells in RIPA buffer was prepared as for TG contents above. Western blot was performed as described previously.^62^ Signal was captured by enhanced chemiluminescence using iBright (ThermoFisher). Rabbit monoclonal LC3A/B antibody (#12741, Cell signaling) was used at 1:1000.

### Bodipy C12 secretion

Bodipy C 12 secretion from PLIN2 KD INS-1 cells was measured following a published protocol.^41^ In brief, one day after transfection, 2 μM Bodipy C12 was added to INS-1 cells cultured in INS-1 medium to metabolically label the cells overnight. Then, medium was removed and changed to INS-1 medium supplemented with 1% FA-free BSA and 0.5 μM OA for 4 h at 37 °C in 5% CO_2_ to capture Bodipy C12 secreted into medium by FA-free BSA. Low concentration of OA was added to prevent extreme deprivation of FA while leaving a sufficient capacity to capture FA by FA-free BSA. At the end of 4 h, medium was saved, and cells were lysed in RIPA buffer. Concentration of Bodipy C12 in medium and RIPA buffer was measured using known concentration of Bodipy C12 as standard and by recording fluorescent emission at 575 nm after 545 nm excitation using a microplate reader.

### Measurement of lysosome-mitochondria contact

In Cont and PLIN2 KD INS1 cells, lysosomes were labeled by transducing cells with CellLight lysosome-GFP (ThermoFisher) and mitochondria were marked by MitoTracker Deep Red (ThermoFisher) following the manufacture’s instruction. The cells were fixed and imaged by Zeiss LSM 980 Airyscan confocal microscope. The contacts between mitochondria and lysosomes are quantitated by a using pre-programmed macro script in ImageJ/Fiji (1.53v, NIH), which was modified from ContactJ macro script.^66^

### Statistics

Data are presented as mean ± SEM unless otherwise stated in the figure caption. Differences of numeric parameters between two groups were assessed with Student’s t-tests. Pairwise comparison was used for values obtained in human islets from the same donor. Welch correction was applied when variances between two groups were significantly different by F test using Prism 8 (GraphPad, La Jolla, CA). Multiple group comparisons used one-way ANOVA with post hoc as indicated. A p < 0.05 was considered significant.

## Acknowledgement

YI is supported by the National Institutes of Health (R01-DK090490), Department of Veteran Affairs (I01 BX005107), and Fraternal Orders of Eagles Diabetes Research Center at the University of Iowa. MG was supported by T32 CA078586 (Free Radical and Radiation Biology, University of Iowa) from NIH. TT is supported by R35 GM144057 from NIH. A Zeiss 980 confocal microscope located in the University of Iowa Central Microscopy Research Facility (CMRF) was funded by the Roy J Carver Charitable Trust. A part of human pancreatic islets was provided by the NIDDK-funded Integrated Islet Distribution Program (IIDP) (RRID:SCR _014387) at City of Hope, NIH Grant # U24DK098085. Human islets were also provided by the Alberta Diabetes Institute Islet Core at the University of Alberta in Edmonton with the assistance of the Human Organ Procurement and Exchange (HOPE) program, Trillium Gift of Life Network (TGLN), and other Canadian organ procurement organizations.

## REFERENCES

1. Halban, P.A., et al. beta-cell failure in type 2 diabetes: postulated mechanisms and prospects for prevention and treatment. J Clin Endocrinol Metab 99, 1983–1992 (2014).

2. Imai Y, E.-L.D., Peachee S J. Pancreatic islet adaptation and failure in metabolic syndrome. in Metabolic Syndrome, Comprehensive Textbook (ed. AhimA, R.) 385–404 (Springer Nature, 2024).

3. Imai, Y., Dobrian, A.D., Morris, M.A. & Nadler, J.L. Islet inflammation: a unifying target for diabetes treatment? Trends Endocrinol Metab 24, 351–360 (2013).

4. Miller, M.R., et al. Levels of free fatty acids (FFA) are associated with insulin resistance but do not explain the relationship between adiposity and insulin resistance in Hispanic Americans: the IRAS Family Study. The Journal of clinical endocrinology and metabolism 97, 3285–3291 (2012).

5. Morita, S., Shimajiri, Y., Sakagashira, S., Furuta, M. & Sanke, T. Effect of exposure to non-esterified fatty acid on progressive deterioration of insulin secretion in patients with Type 2 diabetes: a long-term follow-up study. Diabet Med 29, 980–985 (2012).

6. Morze, J., et al. Metabolomics and Type 2 Diabetes Risk: An Updated Systematic Review and Meta-analysis of Prospective Cohort Studies. Diabetes Care 45, 1013–1024 (2022).

7. Taylor, R., et al. Remission of Human Type 2 Diabetes Requires Decrease in Liver and Pancreas Fat Content but Is Dependent upon Capacity for beta Cell Recovery. Cell Metab 28, 547–556 e543 (2018).

8. Wagner, R., et al. Metabolic implications of pancreatic fat accumulation. Nat Rev Endocrinol 18, 43–54 (2022).

9. Imai, Y., Cousins, R.S., Liu, S., Phelps, B.M. & Promes, J.A. Connecting pancreatic islet lipid metabolism with insulin secretion and the development of type 2 diabetes. Ann N Y Acad Sci 1461, 53–72 (2020).

10. Listenberger, L.L., et al. Triglyceride accumulation protects against fatty acid-induced lipotoxicity. Proc Natl Acad Sci U S A 100, 3077–3082 (2003).

11. Piccolis, M., et al. Probing the Global Cellular Responses to Lipotoxicity Caused by Saturated Fatty Acids. Mol Cell 74, 32–44 e38 (2019).

12. Mathiowetz, A.J. & Olzmann, J.A. Lipid droplets and cellular lipid flux. Nat Cell Biol 26, 331–345 (2024).

13. Zadoorian, A., Du, X. & Yang, H. Lipid droplet biogenesis and functions in health and disease. Nat Rev Endocrinol 19, 443–459 (2023).

14. Najt, C.P., Devarajan, M. & Mashek, D.G. Perilipins at a glance. J Cell Sci 135, jcs259501 (2022).

15. Sztalryd, C. & Brasaemle, D.L. The perilipin family of lipid droplet proteins: Gatekeepers of intracellular lipolysis. Biochim Biophys Acta Mol Cell Biol Lipids 1862, 1221–1232 (2017).

16. Faleck, D.M., et al. Adipose differentiation-related protein regulates lipids and insulin in pancreatic islets. Am J Physiol Endocrinol Metab 299, E249–257 (2010).

17. Chen, E., et al. PLIN2 is a Key Regulator of the Unfolded Protein Response and Endoplasmic Reticulum Stress Resolution in Pancreatic beta Cells. Sci Rep 7, 40855 (2017).

18. Mishra, A., et al. Perilipin 2 downregulation in beta cells impairs insulin secretion under nutritional stress and damages mitochondria. JCI Insight 6, e144341 (2021).

19. Tong, X. & Stein, R. Lipid Droplets Protect Human beta-Cells From Lipotoxicity-Induced Stress and Cell Identity Changes. Diabetes 70, 2595–2607 (2021).

20. Rambold, A.S., Cohen, S. & Lippincott-Schwartz, J. Fatty acid trafficking in starved cells: regulation by lipid droplet lipolysis, autophagy, and mitochondrial fusion dynamics. Dev Cell 32, 678–692 (2015).

21. Benner, C., et al. The transcriptional landscape of mouse beta cells compared to human beta cells reveals notable species differences in long non-coding RNA and protein-coding gene expression. BMC Genomics 15, 620 (2014).

22. Phelps, E.A., et al. Advances in pancreatic islet monolayer culture on glass surfaces enable super-resolution microscopy and insights into beta cell ciliogenesis and proliferation. Sci Rep 7, 45961 (2017).

23. Brennecke, B.R., et al. Utilization of commercial collagens for preparing well-differentiated human beta cells for confocal microscopy. Front Endocrinol (Lausanne) 14, 1187216 (2023).

24. Wolins, N.E., et al. S3-12, Adipophilin, and TIP47 package lipid in adipocytes. J Biol Chem 280, 19146–19155 (2005).

25. Sztalryd, C., et al. Functional compensation for adipose differentiation-related protein (ADFP) by Tip47 in an ADFP null embryonic cell line. J Biol Chem 281, 34341–34348 (2006).

26. Stiles, L. & Shirihai, O.S. Mitochondrial dynamics and morphology in beta-cells. Best Pract Res Clin Endocrinol Metab 26, 725–738 (2012).

27. Bosch, M., Parton, R.G. & Pol, A. Lipid droplets, bioenergetic fluxes, and metabolic flexibility. Semin Cell Dev Biol 108, 33–46 (2020).

28. Grabner, G.F., Xie, H., Schweiger, M. & Zechner, R. Lipolysis: cellular mechanisms for lipid mobilization from fat stores. Nat Metab 3, 1445–1465 (2021).

29. Liu, S., et al. Adipose Triglyceride Lipase Is a Key Lipase for the Mobilization of Lipid Droplets in Human beta-Cells and Critical for the Maintenance of Syntaxin 1a Levels in beta-Cells. Diabetes 69, 1178–1192 (2020).

30. Pearson, G.L., et al. Lysosomal acid lipase and lipophagy are constitutive negative regulators of glucose-stimulated insulin secretion from pancreatic beta cells. Diabetologia 57, 129–139 (2014).

31. Stillman, J.M., et al. Lipofuscin-like autofluorescence within microglia and its impact on studying microglial engulfment. Nat Commun 14, 7060 (2023).

32. Tong, X., Liu, S., Stein, R. & Imai, Y. Lipid Droplets’ Role in the Regulation of beta-Cell Function and beta-Cell Demise in Type 2 Diabetes. Endocrinology 163, bqac007 (2022).

33. Singh, R., et al. Autophagy regulates lipid metabolism. Nature 458, 1131–1135 (2009).

34. Schulze, R.J., et al. Direct lysosome-based autophagy of lipid droplets in hepatocytes. Proc Natl Acad Sci U S A 117, 32443–32452 (2020).

35. Cali, T. & Brini, M. Quantification of organelle contact sites by split-GFP-based contact site sensors (SPLICS) in living cells. Nat Protoc 16, 5287–5308 (2021).

36. Chung, J., et al. The Troyer syndrome protein spartin mediates selective autophagy of lipid droplets. Nat Cell Biol 25, 1101–1110 (2023).

37. Abu-Remaileh, M., et al. Lysosomal metabolomics reveals V-ATPase- and mTOR-dependent regulation of amino acid efflux from lysosomes. Science 358, 807–813 (2017).

38. Tan, A., Prasad, R., Lee, C. & Jho, E.H. Past, present, and future perspectives of transcription factor EB (TFEB): mechanisms of regulation and association with disease. Cell Death Differ 29, 1433–1449 (2022).

39. Petiot, A., Ogier-Denis, E., Blommaart, E.F., Meijer, A.J. & Codogno, P. Distinct classes of phosphatidylinositol 3’-kinases are involved in signaling pathways that control macroautophagy in HT-29 cells. J Biol Chem 275, 992–998 (2000).

40. Kim, Y.C. & Guan, K.L. mTOR: a pharmacologic target for autophagy regulation. J Clin Invest 125, 25–32 (2015).

41. Cui, W., et al. Lipophagy-derived fatty acids undergo extracellular efflux via lysosomal exocytosis. Autophagy 17, 690–705 (2021).

42. Eizirik, D.L., Korbutt, G.S. & Hellerstrom, C. Prolonged exposure of human pancreatic islets to high glucose concentrations in vitro impairs the beta-cell function. J Clin Invest 90, 1263–1268 (1992).

43. Cheruiyot, A., et al. Sustained hyperglycemia specifically targets translation of mRNAs for insulin secretion. J Clin Invest 134(2023).

44. Taddeo, E.P., et al. Mitochondrial Proton Leak Regulated by Cyclophilin D Elevates Insulin Secretion in Islets at Nonstimulatory Glucose Levels. Diabetes 69, 131–145 (2020).

45. Corkey, B.E., Deeney, J.T. & Merrins, M.J. What Regulates Basal Insulin Secretion and Causes Hyperinsulinemia? Diabetes 70, 2174–2182 (2021).

46. Kaushik, S. & Cuervo, A.M. Degradation of lipid droplet-associated proteins by chaperone-mediated autophagy facilitates lipolysis. Nat Cell Biol 17, 759–770 (2015).

47. Tsai, T.H., et al. The constitutive lipid droplet protein PLIN2 regulates autophagy in liver. Autophagy, 1–15 (2017).

48. Griffin, J.D., Bejarano, E., Wang, X.D. & Greenberg, A.S. Integrated Action of Autophagy and Adipose Tissue Triglyceride Lipase Ameliorates Diet-Induced Hepatic Steatosis in Liver-Specific PLIN2 Knockout Mice. Cells 10(2021).

49. Wu, X., et al. Lipid Droplets Maintain Energy Homeostasis and Glioblastoma Growth via Autophagic Release of Stored Fatty Acids. iScience 23, 101569 (2020).

50. Masuda, Y., et al. ADRP/adipophilin is degraded through the proteasome-dependent pathway during regression of lipid-storing cells. J Lipid Res 47, 87–98 (2006).

51. Gerber, P.A. & Rutter, G.A. The Role of Oxidative Stress and Hypoxia in Pancreatic Beta-Cell Dysfunction in Diabetes Mellitus. Antioxid Redox Signal 26, 501–518 (2017).

52. Ngo, J., et al. Mitochondrial morphology controls fatty acid utilization by changing CPT1 sensitivity to malonyl-CoA. EMBO J 42, e111901 (2023).

53. Schlaepfer, I.R. & Joshi, M. CPT1A-mediated Fat Oxidation, Mechanisms, and Therapeutic Potential. Endocrinology 161(2020).

54. Prentki, M., Matschinsky, F.M. & Madiraju, S.R. Metabolic signaling in fuel-induced insulin secretion. Cell Metab 18, 162–185 (2013).

55. Schott, M.B., et al. Lipid droplet size directs lipolysis and lipophagy catabolism in hepatocytes. J Cell Biol 218, 3320–3335 (2019).

56. Chiariello, A., et al. Downregulation of PLIN2 in human dermal fibroblasts impairs mitochondrial function in an age-dependent fashion and induces cell senescence via GDF15. Aging Cell, e14111 (2024).

57. Najt, C.P., et al. Liver-specific loss of Perilipin 2 alleviates diet-induced hepatic steatosis, inflammation, and fibrosis. Am J Physiol Gastrointest Liver Physiol 310, G726–738 (2016).

58. Bosch, M., Sweet, M.J., Parton, R.G. & Pol, A. Lipid droplets and the host-pathogen dynamic: FATal attraction? J Cell Biol 220(2021).

59. Harata, M., et al. Delivery of shRNA via lentivirus in human pseudoislets provides a model to test dynamic regulation of insulin secretion and gene function in human islets. Physiol Rep 6, e13907 (2018).

60. Singh, B., Khattab, F. & Gilon, P. Glucose inhibits glucagon secretion by decreasing [Ca(2+)](c) and by reducing the efficacy of Ca(2+) on exocytosis via somatostatin-dependent and independent mechanisms. Mol Metab 61, 101495 (2022).

61. Hsieh, K., et al. Perilipin family members preferentially sequester to either triacylglycerol-specific or cholesteryl-ester-specific intracellular lipid storage droplets. J Cell Sci 125, 4067–4076 (2012).

62. Trevino, M.B., et al. Perilipin 5 regulates islet lipid metabolism and insulin secretion in a cAMP-dependent manner: implication of its role in the postprandial insulin secretion. Diabetes 64, 1299–1310 (2015).

63. Merrill, R.A., Flippo, K.H., Strack, S. Measuring Mitochondrial Shape with ImageJ. in Neuromethods, Techniques to Investigate Mitochondrial Function in Neurons, Vol. 123 (ed. Strack, S., Usachev, Y. M.) 31–48 (Springer, U.S.A., 2017).

64. Slupe, A.M., et al. A calcineurin docking motif (LXVP) in dynamin-related protein 1 contributes to mitochondrial fragmentation and ischemic neuronal injury. J Biol Chem 288, 12353–12365 (2013).

65. Yim, W.W., Yamamoto, H. & Mizushima, N. Annexins A1 and A2 are recruited to larger lysosomal injuries independently of ESCRTs to promote repair. FEBS Lett 596, 991–1003 (2022).

66. Martin, G., et al. ContactJ: Characterization of lipid droplet-mitochondrial contacts using fluorescence microscopy and image analysis. F1000Res 10, 263 (2021).

