## Supplementary Figures for "Lipid droplet protein Perilipin 2 is critical for the regulation of insulin secretion through beta cell lipophagy and glucagon expression in pancreatic islets"

**Supplementary methods**

***Validation of SPLICS***

Designing of SPLICS-P2A-LD–LYSO is described in Methods. To validate targeting of $GFP_{1-10}$ fragment to the surface of lipid droplets, INS-1 cells were transduced with SPLICS-P2A-LD–LYSO lentivirus for 3 days, fixed, and immunostained by anti-GFP antibody (#2925, Cell Signaling Technology) that recognizes both a full GFP and $GFP_{1-10}$ fragment.[1] To validate the proper targeting of GFP $\beta_{11}$ fragment at lysosome surface, $GFP_{1-10}$ was expressed in cytosol by transiently transfecting pcDNA3.1-GFP (1-10) (Addgene #70219, a gift from Dr. Bo Huang)[2] in Hela cells that were transduced with SPLICS-P2A-LD–LYSO lentivirus. 3 days after transfection, cells were fixed. Then, GFP was immunostained by anti-GFP antibody (#2925, Cell Signaling Technology) and lysosomes were visualized by LysoTracker Deep Red (ThermoFisher). All the Imaging were performed by Zeiss LSM 980 Airyscan confocal microscope and analyzed via ImageJ/Fiji (1.53v, NIH).

### References


1. Cali, T. & Brini, M. Quantification of organelle contact sites by split-GFP-based contact site sensors (SPLICS) in living cells. *Nat Protoc* **16**, 5287-5308 (2021).
2. Kamiyama, D.*, et al.* Versatile protein tagging in cells with split fluorescent protein. *Nat Commun* **7**, 11046 (2016).

**Supplementary Figure 1.**

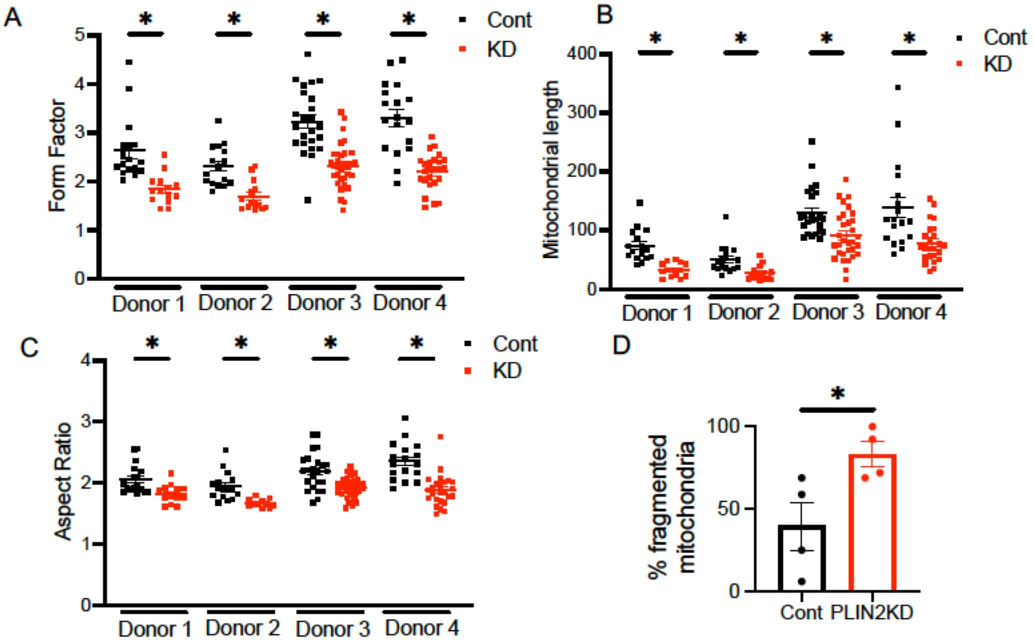


The morphology of mitochondria was assessed in beta cells marked by anti-insulin antibody in four human donor islets. (A) Form factors, (B) mitochondrial length, and (C) aspect ratio analyzed in n=13 to 32 images per condition. (D) % of fragmented mitochondria was defined for each donor as aspect ratio below 2 in (C), n=4 donors. Data are mean ± SEM. *; $p<0.05$ by Student's t-test.

**Supplementary Figure 2**

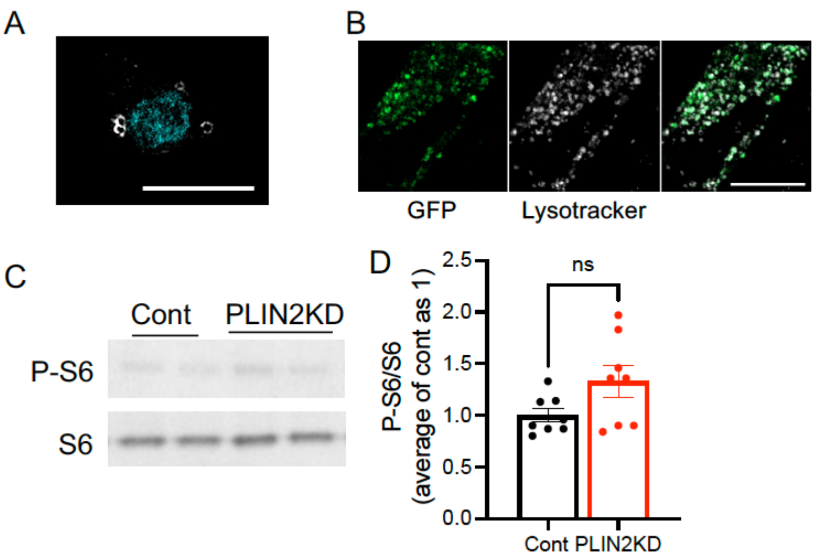


(A) INS-1 cells expressing lenti-SPLICS-P2A-LD–LYSO were incubated with anti-GFP antibody to visualize GFP 1-10 showing typical ring-like distribution of a LD surface protein (white). Nucleus marked by DAPI is shown in cyan. (B) Hela cells were transduced by lenti-SPLICS-P2A-LD–LYSO and pcDNA3.1-GFP (1-10) (Addgene #70219, a gift from Dr. Bo Huang)[2] so that the distribution of $\beta_{11}$ can be visualized as green fluorescence.[1] The distribution of GFP and lysosome visualized by lysotracker showed significant overlap. Scale bars are 10 µm. (C-D) Western blot assessed Phospho-S6 ribosomal protein (P-S6, #5364 from Cell Signaling at 1:1000) signal corrected for total S6 Ribosomal protein (S6, #2217 from Cell Signaling at 1:1000) in Cont and PLIN2KD INS-1 cells. (C) Representative blot and (D) the ratio of P-S6/S6 from densitometry obtained from three independent experiments and expressed the average of Cont as 1. n=8.

**Supplementary Figure 3**

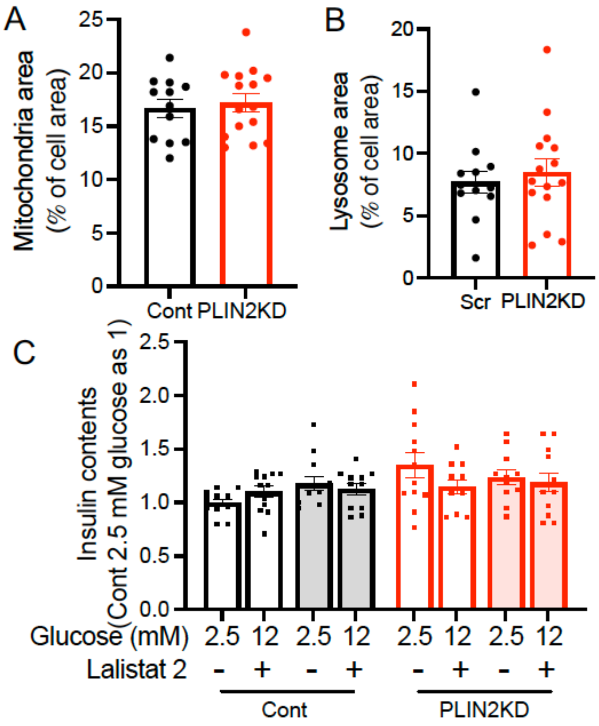


(A) Mitochondria area and (B) lysosome area corrected for cell area in control (Cont) and PLIN2 downregulated (PLIN2KD) INS-1 cells in which lysosomes were marked by CellLigh Lysosomes-GFP (green) and mitochondria by MitoTracker deep red. n= 12 to 15 images. Representative of three independent experiments. (C) Cont and PLIN2 KD INS-1 cells were cultured with or without lalistat 2 (Lali2) for 16 h and insulin contents corrected for protein contents were expressed taking the average of Cont at 2.5 mM glucose as 1. n=12 combining four independent experiments. Data represent mean ± SEM.

**Supplementary Figure 4**

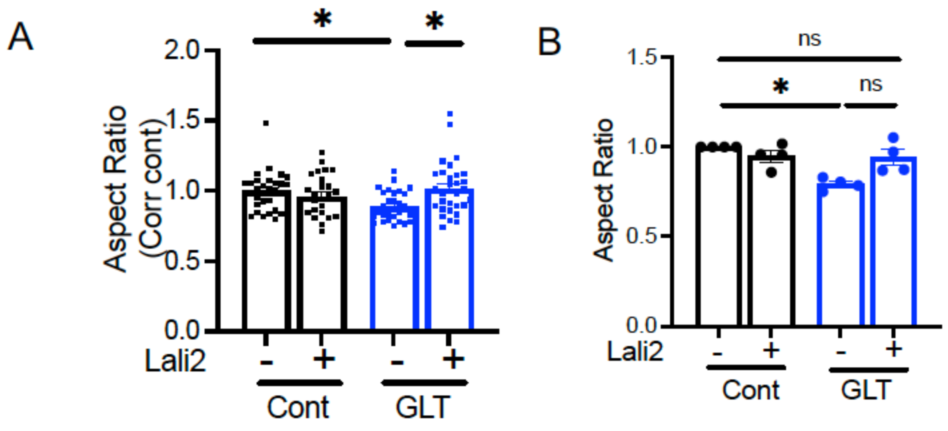


(A) Aspect ratio of mitochondria in INS-1 cells cultured in RPMI medium containing 11.1 mM glucose (Cont) or 20 mM glucose + 0.13 mM OA+ 0.26 mM PA (GLT) in the presence or absence of 20 µM lalistat 2 (Lali2) for overnight in the presence of C12. After fixation, mitochondria were visualized by anti-HSP antibody. n= 28 to 32 images from three independent experiments. (B) Aspect ratio of mitochondria in human beta cells cultured in CMRL1066 medium containing 5.5 mM glucose (Cont) or CMRL1066 medium at 20 mM glucose + 0.16 mM OA+ 0.32 mM PA (GLT) with or without 10 µM Lali2 overnight. Mitochondria were visualized by CellLight Mitochondria-GFP. Beta cells were identified by insulin antibody. Each dot represents the average for each donor sample, n= 4 donors. Data represent mean ± SEM.
